# The structures of tetrameric avian immunoglobulin A are uniquely structured to support mucosal immunity

**DOI:** 10.1101/2025.09.19.677352

**Authors:** Rebecca M. Schneider, Qianqiao Liu, Beth M. Stadtmueller

## Abstract

Birds and mammals assemble polymeric (p) forms of immunoglobulin (Ig) A, which are transported to mucosal surfaces and released as secretory (S) IgA, the predominant mucosal antibody. Mammalian SIgA, which is typically dimeric, is well characterized; however, avian IgA structure-function relationships and how they compare to those in mammals remains unclear. Here we report the cryo-electron microscopy structures of mallard duck pIgA and SIgA at 3.76-Å and 3.21-Å resolution, respectively. Both structures revealed four IgA monomers linked through one joining chain (JC) and adopted a non-planar conformation; a heavy chain domain absent in mammals, extended the tetramer’s non-planar geometry toward antigen binding sites. The SIgA structure revealed four secretory component (SC) domains binding IgA and JC analogous to four of the five mammalian SC domains; however, SC and JC N-terminal extensions, absent in mammals, mediated additional interfaces between complex components. Experiments comparing mammalian and avian SC binding to cognate and non-cognate IgA ligands demonstrated that species-specific structural differences correlated with differences in SC binding and that adding an N-terminal extension to mammalian SC enhanced binding to avian pIgA. Together, results and comparison to published structures indicate that avian IgA components co-evolved unique elements compared to mammalian IgA, including a predominantly tetrameric state, and suggest that birds may employ different IgA effector mechanisms to combat mucosal pathogens compared to mammals. These findings are broadly relevant to understanding IgA evolution and function and combating zoonotic disease such as avian influenza A virus.

**One Sentence Summary:** Cryo-electron microscopy structures together with biochemical experiments demonstrate similarities and differences between mammalian and avian secretory immunoglobulin A relevant to understanding differences in immune response to common pathogens such as influenza virus.

## INTRODUCTION

The vertebrate mucosa forms a critical protective barrier against external threats. In mammals and birds, secretory (S) immunoglobulin (Ig) A (SIgA) populates this barrier, serving as the predominant mucosal antibody and protecting the host from bacterial and viral pathogens (*1–3*). To fulfill species-specific immune needs, SIgA has evolved unique polymeric (p) structures and effector functions distinct from monomeric IgA and other circulatory antibodies (*4, 5*). Additionally, variation in SIgA structures and effector functions has been associated with species, isoform, and allotype, and has likely been driven in part by host-pathogen co-evolution (*6*). While our understanding of mammalian SIgA structure-function relationships continues to grow, knowledge of avian SIgA remains limited. Given that birds experienced distinct evolutionary history compared to mammals and are recognized as reservoirs for zoonotic diseases, such as those caused by influenza A viruses (IAV) (*7–9*), understanding how avian SIgA differs from mammalian SIgA and how it protects birds from pathogens may offer critical insights to prevent disease spillover to humans.

In mammals, SIgA production begins in IgA-expressing plasma cells, which assemble pIgA by combining two to five IgA monomers with one joining chain (JC) (*10–13*). Each IgA monomer (mIgA) comprises two heavy chains (HCs) and two light chains (LCs). Each HC contains a variable domain (V_H_), three Ig constant domains (Cα1, Cα2, Cα3), and an eighteen-residue C-terminal tailpiece (Tp), while each LC includes one variable domain (V_L_) and one constant domain (C_L_). The monomer includes two antigen-binding fragments (Fabs) and one fragment crystallizable (Fcα), with the latter conferring effector function specificity and mediating critical contacts with the JC and adjacent mIgAs in pIgA complexes. The cryo-electron microscopy (cryo-EM) structure of mouse dimeric (d) IgA (*11*) (containing two mIgA and one JC) revealed that the JC N-terminal half (termed the JC_core_) forms a β-sandwich-like assembly with the HC Tps and uses two β-hairpins (C-terminal half) to contact Cα3 domains in both Fcαs (*10*). The resulting complex is asymmetric, with the two mIgAs bent and tilted relative to each other (*11*). This dIgA structure, which is expected to represent the predominant form of IgA secreted by plasma cells, is subsequently bound by the polymeric Ig receptor (pIgR) on the basolateral side of epithelial cells. The pIgR contains five Ig-like domains (D1-D5) with loops structurally similar to antibody complementarity-determining regions (CDRs) and transcytoses pIgA to the apical side of cells where it is proteolytically cleaved, releasing SIgA, a complex of the cleaved pIgR ectodomain (secretory component; SC) and pIgA, into the mucosal lumen (*14*). Cryo-EM structures of human and mouse dimeric SIgA (*10, 11*) revealed that the SC binds to one face of dIgA, contacting both Fcαs, the Tps, and the JC, thereby increasing the asymmetry of the complex while leaving the overall conformation of dIgA similar to its unliganded structure (*1*).

In mucosal secretions, SIgA structure supports unique effector functions typically dominated by biophysical mechanisms such as antigen crosslinking and subsequent removal through peristalsis (e.g., in the gut) or the mucociliary escalator (e.g., in the airways) (*1*). Modeling of Fabs onto dimeric SIgA structures revealed that that the geometric relationships between the two mIgAs influence the positions that Fabs can occupy and thus are likely to influence antigen binding and cross-linking mechanisms (*1, 11*). SIgA effector mechanisms can also involve IgA-specific Fc Receptors (FcαRs) expressed on host cell surfaces. Upon binding IgA-immune complexes, these receptors can trigger processes such as phagocytosis or respiratory burst (*15–19*). Two accessible FcαR binding sites have been identified in mammalian dimeric SIgA structures, yet complex asymmetry may limit binding at one site in some cases (*16*). While over 90% of SIgA in humans and mice is dimeric, approximately 10% exists as higher-order polymers (i.e., tetramers or pentamers) (*1, 20*). Published cryo-EM structures revealed the cores of human tetrameric and pentameric SIgA (*11*) to be largely superimposable with those of dimeric SIgA (*10, 11*), with the addition of two or three mIgAs that add Tps to the central β-sandwich-like assembly but do not interact with the JC and SC (*1*). Together, mammalian IgA structures and Fab modeling suggest distinct geometric relationships between IgA Fabs and FcαR binding site accessibility in higher-order polymeric SIgAs compared to mIgA or dIgA. Therefore, factors such as polymeric state, antigen identity, FcαR identity, and host immune cell activity are thought to influence IgA effector functions (*1, 13*).

The impact of SIgA’s structural features on host responses to specific pathogens remains largely unclear, yet crucial to understanding and preventing disease. Investigating how SIgA protects against zoonotic diseases originating in avian hosts is particularly important, as these hosts have evolved SIgAs that are thought to differ from mammalian variants and remain largely uncharacterized. Avian IgA HCs contain four Ig constant domains (Cα1, Cα2, Cα3, and Cα4) rather than three (*21, 22*) and molecular weight measurements of endogenous avian IgA suggested a predominantly tetrameric rather than dimeric structure (*23*). Additionally, the avian pIgR ectodomain (SC) contains four Ig-like domains, lacking a domain homologous to mammalian pIgR D2 (*24*). To further understand the evolution of vertebrate IgA structure and effector functions, as well as to investigate differences between birds and mammals relevant to understanding zoonotic diseases, we targeted the structures of avian pIgA and SIgA, specifically those expressed by mallard ducks (*Anas platyrhynchos*). Serving as a reservoir for AIV, mallard ducks typically avoid serious illness while transmitting fatal disease to other birds and mammals (*7–9, 25*). We determined pIgA and SIgA cryo-EM structures, lacking Fabs, at 3.76-Å and 3.21-Å resolution, respectively. Comparative analysis with mammalian structures, as well as supporting biochemical experiments, revealed unique structural and geometric features that likely translate into differences in antigen engagement and effector functions compared to mammals.

## RESULTS

### The cryo-EM structure of avian IgA

To investigate the structures of avian IgA we co-expressed mallard duck HC (containing Cα2-Cα3-Cα4-tp) and JC constructs and determined the cryo-EM structure, hereafter referred to as (*Anas platyrhynchos*; *ap*) JFcα to an average resolution of 3.79 Å (Fig. 1A; fig. S1, A to E; table S1). Local resolution was variable (fig. S1F); however, density for most main chain and side chain atoms in Cα4, Tp, and JC residues were well-resolved, along with those in Cα3 participating in molecular interfaces (fig. S1, G to I). Seven N-terminal residues and the two C-terminal residues (137–138) of the JC were disordered, along with a subset of residues in the eight HC C-terminal Tps (residues 423-440). Local resolution among Cα3 domain residues varied from 3.3 Å to 8.0 Å. Main chain and side chain atoms for less resolved regions of Cα3 were built in geometrically reasonable conformations. Local resolution for Cα2 domains varied from 6.5 Å to 11 Å and therefore Cα2 residues were not built; this is consistent with human IgM structures in which Cμ2 is reportedly flexible and poorly resolved (*26, 27*).

**Fig. 1.**
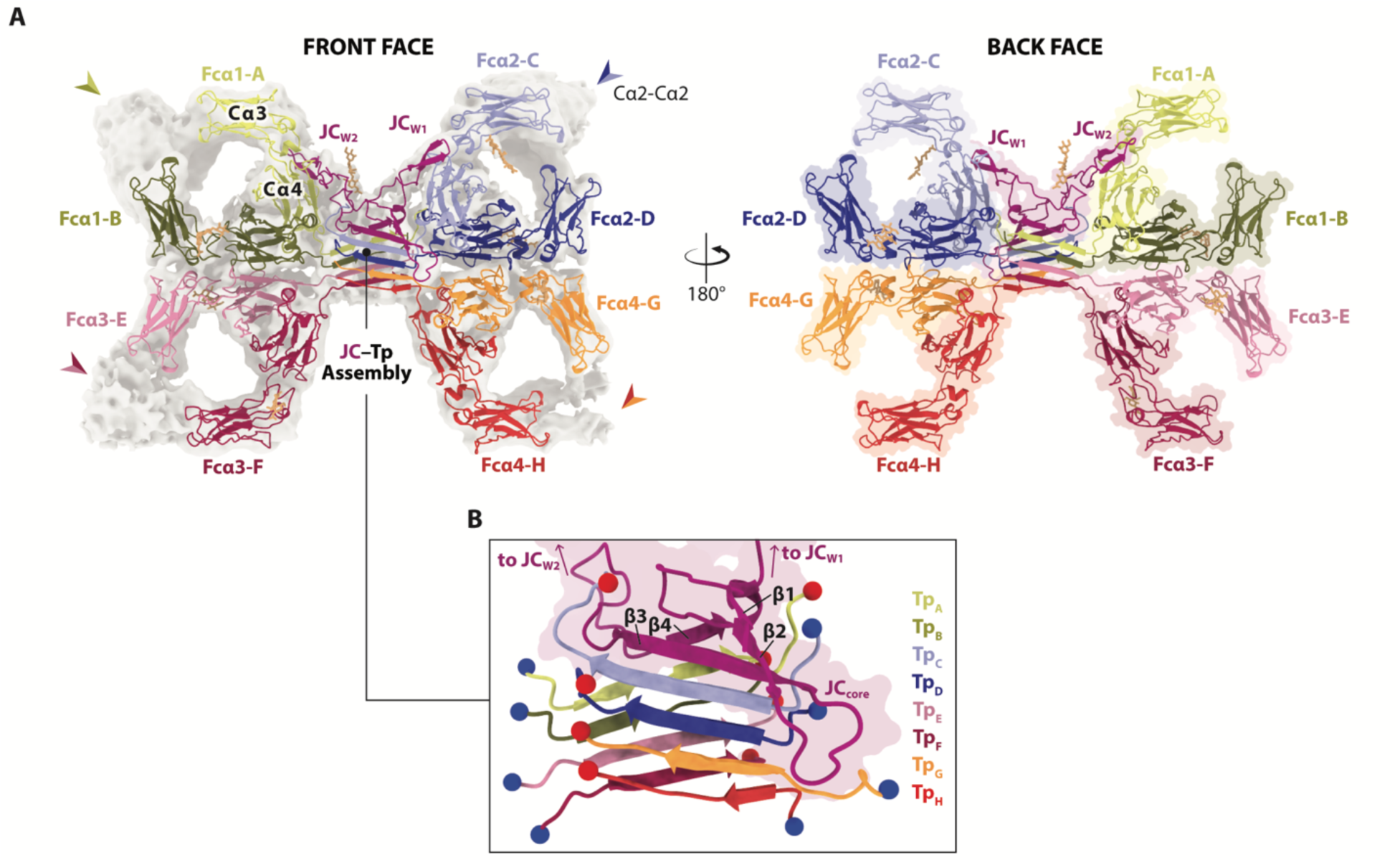
The structure of *ap* JFcα. (**A**) Front and back face views of *ap* JFcα cartoons with glycans shown as sticks; heavy chains (A–H), Fcα subunits (Fcα1–Fcα4), and JC are uniquely colored and labeled. Front face view includes density map with Cα2 domains indicated by arrows. (**B**) Front face cartoon of JC-Tp assembly with JC_core_ shown as a flat molecular surface; chains are colored as in (A) and labeled along with the four JC_core_ β strands. Tp N- and C-termini are indicated by blue and red spheres, respectively.

The refined *ap* JFcα structure revealed a tetrameric IgA with four Fc subunits (Fcα1 to Fcα4) comprising HCs A-H and JC (Fig. 1A). The Fcα subunits are arranged such that each subunit forms an interface with an adjacent subunit, resulting in two Fc-Fc interfaces: the Fcα1-Fcα3 interface and the Fcα2-Fcα4 interface. The center of the complex is occupied by the eight HC tailpieces (Tp_A_ to Tp_G_) which form a β-sandwich-like domain that is further extended by the JC_core_ as JC strands β1, β2, and β3 extend the Tp β-sandwich on the front face (via contacts between β3 and Tp_C_) and β4 strand extends the Tp β-sandwich on the back face (via contacts between β4 strand and Tp_A_) (Fig. 1B). While the β-sheet interactions formed in this JC-Tp assembly dominate the JC_core_-IgA interface, the JC_core_ also makes contacts with residues outside of the Tp region in Fcα1, Fcα2, and Fcα4 (fig. S2, A to C). The second half of the JC sequence is folded into two β-hairpin ‘wings’ (JC_W1_ and JC_W2_); JC_W1_ extends from the center of the complex toward the front face to form contacts with Fcα2 chain C (Fcα2-C) and JC_W2_ extends from the center of the complex toward the back face to contact Fcα1-A (Fig. 1A; fig. S2, A, D and E). Overall, the JC-Tp assembly is structurally similar to those observed in mammalian dIgA and SIgA structures, with an RMSD of 0.741 Å when aligned to Cα atoms in the mouse dIgA JC-Tp assembly and RMSD of 1.602 Å when aligned to Cα atoms in the human tetrameric IgA structure. Contacts with Fcs are also similar between species, although some side chain interactions differ among the structures. Furthermore, we observed that the linker connecting JC_W1_ and JC_W2,_ which is four residues shorter than mammalian counterparts, is ordered in the avian structure whereas a subset of these residues are disordered in mammalian IgA structures. We also observed ordered glycans at seven sites of the 33 potential N-linked glycosylation sites (PNGS) on *ap* JFcα, one on each Fcα1-B, Fcα2-C, Fcα2-D, Fcα3-E, Fcα3-F, Fcα4-G Asn230 and one on JC Asn53, and were able to build two to three bases at each of these sites. Although we were unable to assign additional bases, the map density suggested that HC Asn230 glycans extend beyond 3 bases and may contact other HC residues (fig. S3).

Inspection of the *ap* JFcα structure suggested that the geometric relationships between the four Fcα subunits, relative to each other, were unique (fig. S4A). To quantify these relationships, we defined a central plane (the JFcα plane), which bisects the front face and back face of the Fcα subunits, and then measured the angles between adjacent Fcα subunits lying on the plane (fig. S4, B to D). The central angles between Fcα1-Fcα3 and Fcα2-Fcα4 were ∼61°, whereas the angles between Fcα1-Fcα2 and Fcα3-Fcα4 were ∼119° (Fig. 2A; fig. S4D). Deviations of each Fc from the JFcα central plane were also apparent (Fig. 2B) and inspection of the cryo-EM map suggested that differences in Fcα subunit orientation extend to Cα2 domains (Fig. 2C), which are absent in mammals and may propagate each Fcα subunit’s unique conformational heterogeneity to the associated Fabs. To quantify these relationships, we measured deviations in two directions, termed *twist* and *tilt* (Fig. 2D; fig. S4, E and F). The twist and tilt measurements ranged from −6.1° to +5.9°, except for Fcα2, which displayed a twist of +12.0° and a tilt of −10.9° (Fig. 2D; fig. S4F), demonstrating asymmetry in the complex and suggesting that Fcα2 (and the associated Fabs) can occupy positions more distal from the central plane than other Fcs.

**Fig. 2.**
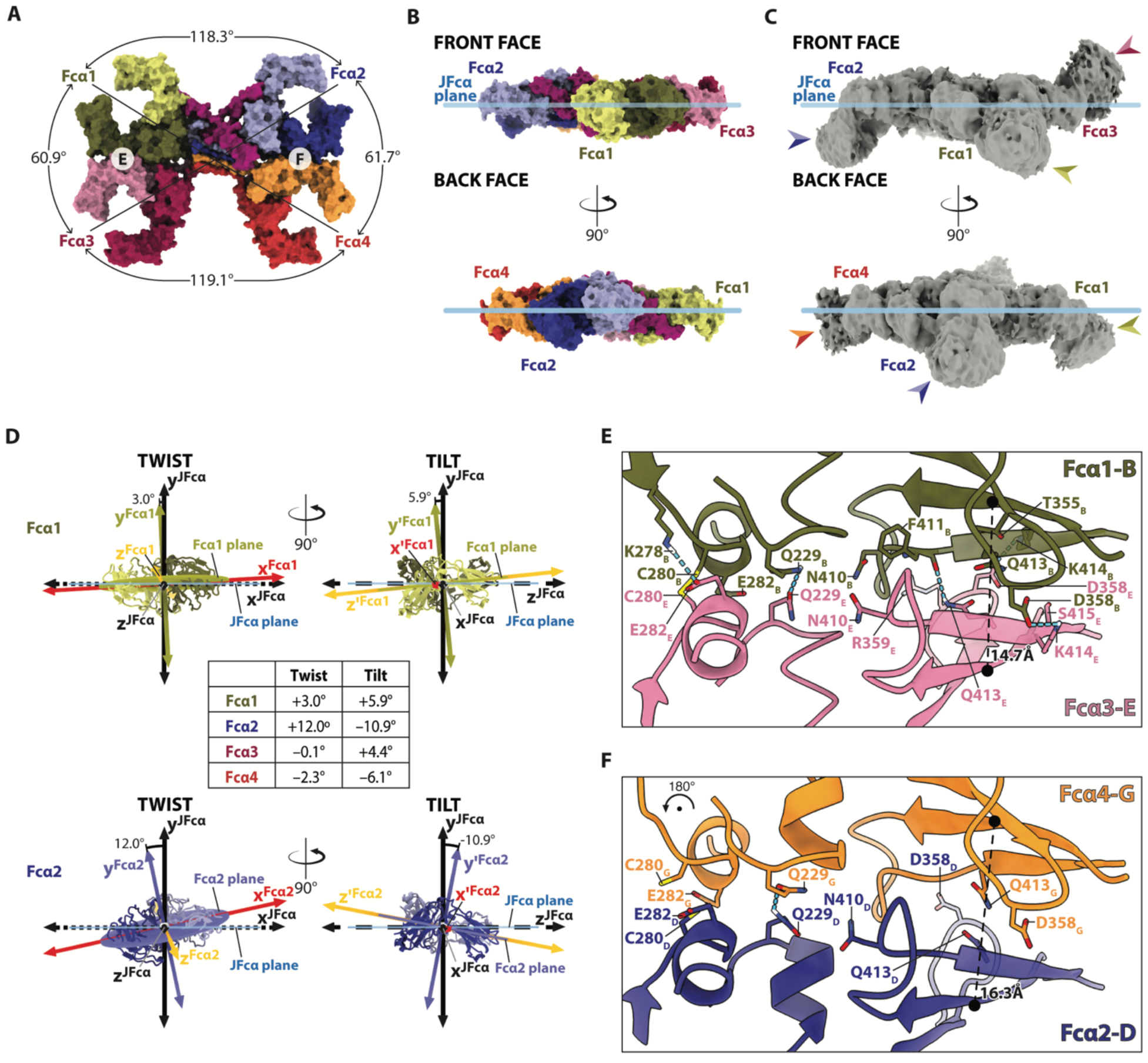
The unique conformation of *ap* JFcα. (**A**) Molecular surface of *ap* JFcα front face with indicated subunits colored as in Fig. 1 and central angles (degrees) between adjacent Fcα subunits labeled. (**B**) Molecular surface of *ap* JFcα side view in three orientations relative to the JFcα plane (light blue); subunits are colored as in (A). (**C**) Cryo-EM density maps shown in the same orientations as (B). Arrows indicate Cα2 domain density. (**D**) Schematic showing twist and tilt measurements for Fcα1 (top) and Fcα2 (bottom) and table of twist and tilt values for Fcα1-Fcα4 (middle). Twist: degrees of displacement from the origin for each subunit plane (colored ellipses) normal (dark olive or dark indigo lines) when rotating about a fixed z-axis (black dashed line; z^JFca^ axis). Tilt: degrees of displacement from the origin for each subunit plane normal after the twist displacement when rotating about a fixed x-axis (black dotted line; x^JFca^ axis) (fig. S4). (**E, F**) Fcα1-Fcα3 (E) and Fcα2-Fcα4 (F) interfaces shown as cartoons; interacting residues shown and labeled. Dashed lines indicate Cys400 distances between respective chains.

Upon identifying geometric differences among Fcα subunits, indicative of complex asymmetry, we inspected molecular interfaces and found differences in contacts between Cα3 and Cα4 domains in the Fcα1-Fcα3 and Fcα2-Fcα4 interfaces, which bury ∼916 Å^2^ and ∼714 Å^2^, respectively. Geometric differences correlated with a higher number of hydrogen bonding, polar, and electrostatic interactions at the Fcα1-Fcα3 interface than the Fcα2-Fcα4 interface (Fig. 2, E and F) and subtle differences in hydrophobic interactions (fig. S5, A to C); the distances separating residue pairs also correlated with the geometric differences (e.g. the distance between Fcα1-B Cys400 and Fcα3-E Cys400 is 14.7 Å, whereas the distance between Fcα2-D Cys400 and Fcα4-G Cys400 is 16.3 Å) (Fig. 2, E and F). Intrachain disulfide bonds are known to stabilize Fc-Fc interfaces in mammalian IgM, which is pentameric, yet are not thought to play a major role in stabilizing mammalian polymeric IgA (*10, 26*). We observe evidence for an inter-Fc disulfide bond between Fcα1-B

Cys280 and Fcα3-E Cys280, adjacent to a hydrogen bond between Fcα1-B Q229 and Fcα3-E Q229 (Fig. 2E; fig. S5D). Although the Fcα2-D Q229 and Fcα4-G Q229 hydrogen bond appears to stabilize the Fcα2-Fcα4 interface, we failed to find evidence for the intrachain disulfide (Fig. 2F; fig. S5E), suggesting that it may not always form.

### The cryo-EM structure of avian SIgA

The *ap* JFcα structure is likely to represent the structure of avian IgA secreted by plasma cells that is subsequently bound by pIgR and released as SIgA. To investigate the structure of avian SIgA, we co-expressed mallard duck HC (containing Cα2-Cα3-Cα4-tp), JC, and pIgR ectodomain (SC; residues 1-458) constructs and determined the cryo-EM structure, *ap* SJFcα, to an average resolution of 3.21 Å (Fig. 3A; fig. S6, A to E; table S1). Local resolution showed similar variability to that observed in the *ap* JFcα structure (fig. S6, F to J); however, local resolution of Fcα1 and Fcα2 Cα4 residues were higher (2.8 Å to 4.2 Å) in the *ap* SJFcα complex. The JC-Tp assembly was also better resolved (2.8 Å to 3.8 Å) along with JC N-terminal residues. Local resolution for SC ranged from 2.8 Å to 5.5 Å with the lowest resolution observed in D4.

**Fig 3.**
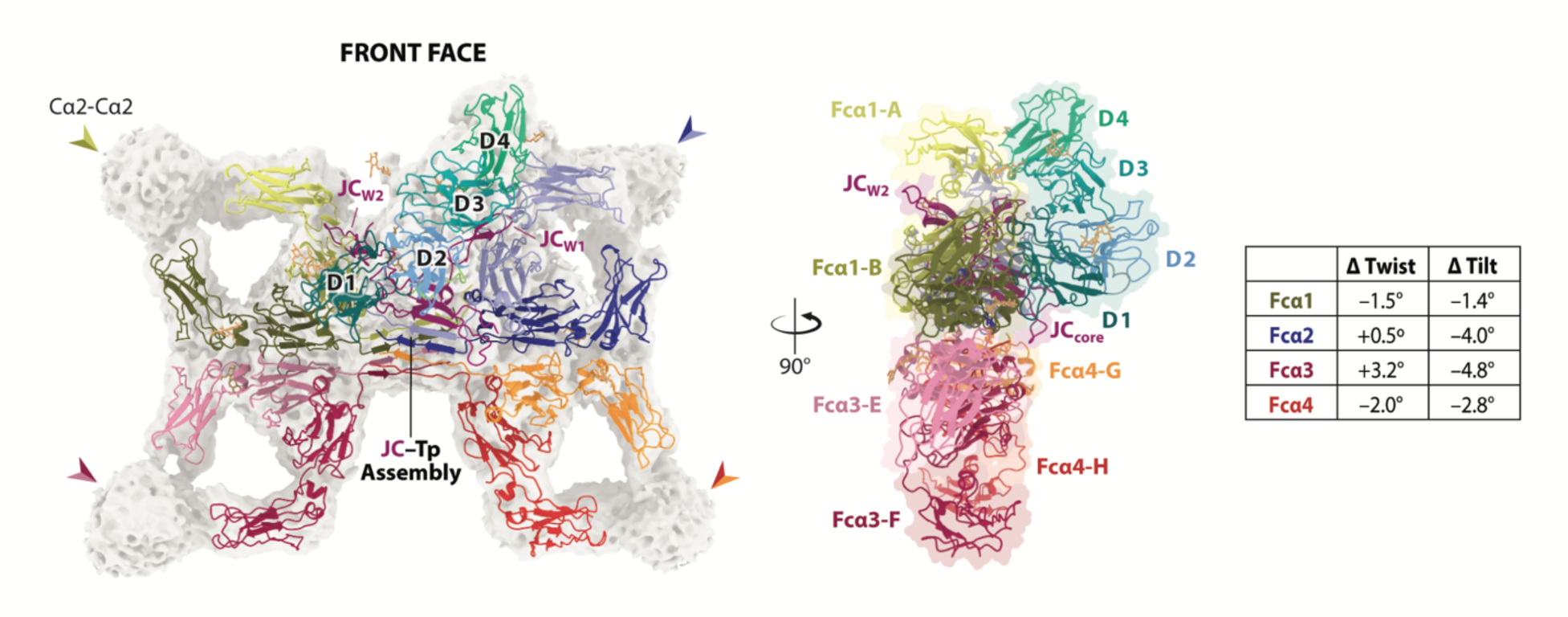
The structure of *ap* SJFcα. Front (left) and side (middle) face views of *ap* SJFcα cartoons with glycans shown as sticks; heavy chains (A-H), Fcα subunits (Fcα1-Fcα4), JC, and SC domains are uniquely colored and labeled. Front face view shows density map with Cα2 domains indicated by arrows. The changes (Δ) in twist and tilt values associated with the addition of SC (i.e. the differences between twist and tilt measured in *ap* JFcα and *ap* SJFcα structures) are reported in the table (right).

The refined *ap* SJFcα structure revealed SC asymmetrically bound to the front face of JFcα with SC domains D1-D4 forming contacts with Fcα1, Fcα2, and the JC. Comparison between *ap* JFcα and *ap* SJFcα structures suggested that SC did not markedly impact the JFcα structure, although some variability was apparent. A global alignment of Cα atoms (with SC removed) revealed an RMSD of 1.603 Å. *Twist* and *tilt* values varied between the two structures (Fig. 3), with the differences in twist (ΔTwist) ranging from −2.0° to +3.2° and the differences in tilt (ΔTilt) ranging from −1.4° to −4.8°. In both complexes, Fcα2 exhibited the largest tilt value; however, the tilt is more negative in *ap* SJFcα (−14.9°) compared to *ap* JFcα (−10.9°). Because a negative tilt value indicates that the subunit is pointed toward the back face of the complex, and in the case of *ap* SJFcα, away from SC, the observed ΔTilt of −4.0° for Fcα2 suggests that SC binding supports and possibly enhances the unique positioning of Fcα2 toward the back face relative to the other subunits. Consistent with these observations, we did not observe marked changes in Fc-Fc interfaces when comparing structures, although evidence for interchain disulfide bonding between Cys280 residues was variable. These observations suggest, that like mammals (*10, 11*), the overall conformation of avian SIgA is stabilized through JC-HC and HC-HC contacts.

### SC_N-ext._ bridges extensive JC-SC interface not observed in mammalian structures

Analysis of the *ap* SJFcα structure and comparison to mammalian SIgA structures revealed both unique and conserved features. For example, in the *ap* SJFcα complex, duck SC buries a larger surface area (∼924 Å^2^) on JC compared to mouse SC (∼597 Å^2^) in the SIgA structure (*11*). This difference is largely due to interfaces that are absent in mammalian SIgA complexes. Avian SC begins with additional N-terminal residues (i.e. eight to eleven residues) that are not conserved in mammals (fig. S7A). In the *ap* SJFcα complex, the 11-residue SC N-terminal extension (SC_N-ext._) was well-resolved; filling a pocket that is unoccupied in mammalian SIgA structures, SC_N-ext._ extends out from D1 and contacts D2, JC, and Fcα2 (Fig. 4, A and B). The first two SC_N-ext._ residues approach D2, positioning residues three to five in contact with both D2, JC, and, to a lesser extent, Fcα2. The subsequent SC_N-ext._ residues (Pro6 to Ile11) bridge interactions between residues in D2 and the JC_core_, which includes JC Arg10 and other residues in the JC β1 strand. Notably, in addition to contacting SC_N-ext._, the JC Arg10 sidechain reaches over the SC_N-ext._ where it is positioned to form a salt bridge with the D2 loop Glu204 side chain, effectively bridging JC interactions with multiple SC motifs (Fig. 4B), whereas in the *ap* JFcα complex, JC Arg10 contacts Fcα2 (fig. S2B). The SC_N-ext._-JC interface is ∼411 Å^2^, which also includes SC_N-ext._ contacts with JC N-terminal residues Asp6, Glu8, and Glu9. We also observed the JC N-terminus hydrogen bonding to JC β2β3 and Fcα2-C Cα4 loop residues, anchoring seven N-terminal JC residues that were disordered in the absence of SC. The N-terminal contact is notable because the first five JC residues, which we term JC_N-ext._, are not conserved in mammalian JC sequences and are divergent among other vertebrates (fig. S7B). Thus, similar to interfaces involving SC_N-ext._, these JC_N-ext._ interfaces are likely unique to ducks and other birds.

**Fig. 4.**
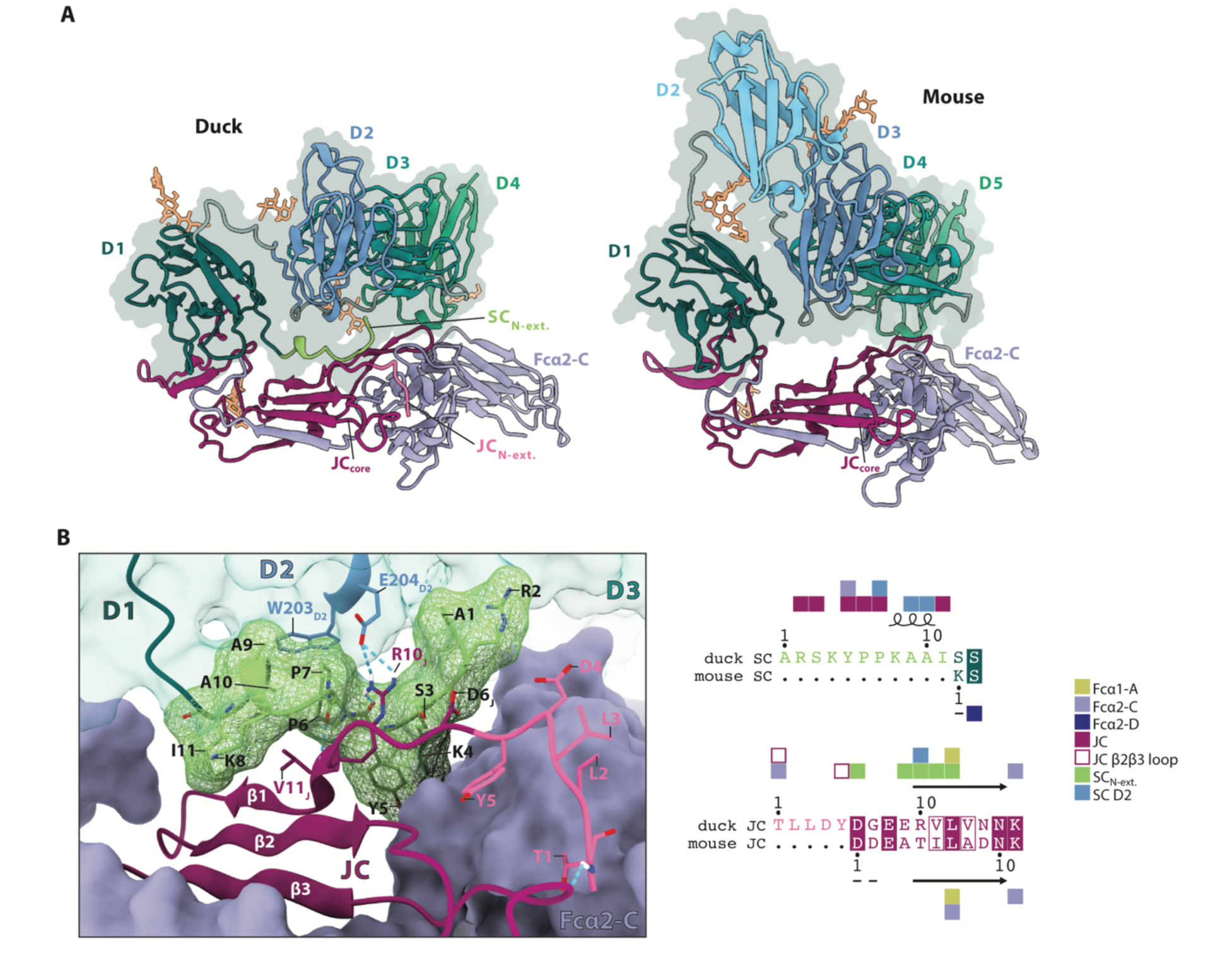
SC and JC N-terminal extensions. (**A**) Cartoon of SC, Fcα2-C, and JC only from *ap* SJFcα (left) and mouse SIgA (PDB: 7JG2, right) structures with glycans shown as sticks. Fcα2-C, SC_N-ext._, SC domains, JC_N-ext._, and JC are uniquely colored and labeled. (**B**) SC_N-ext._ and JC_N-ext._ residues and their contacts in *ap* SJFcα shown as cartoons and sticks. SC domains and JC β strands are labeled. Molecular surface of SC (transparent teal) and Fcα2-C (blue violet) are shown. A mesh molecular surface is shown for the SC_N-ext_. Structure-based sequence alignments of duck and mouse SC and JC N-termini are shown on right with colored squares to indicate contacts according to key. Dashes signify disordered residues.

Together, the SC_N-ext._ and JC_N-ext._ appear to promote a network of stabilizing interfaces that are absent in mammalian SIgA complexes. For example, the binding of SC may drive conformational changes in JC (e.g. the repositioning of Arg10) that facilitate the anchoring of D1 and D2 in SC, as well as the JC_N-ext._. JC N-terminus anchoring promotes more contacts between the JC and Fcα in *ap* SJFcα compared to *ap* JFcα (Fig. 4B; fig. S2B). The resulting larger SC-JC interface in birds, compared to mammals, may provide enhanced molecular stability to support avian SIgA functions. This observation, along with evidence that the SC_N-ext._ is highly conserved in birds (fig. S7, A and B), suggests these N-terminal motifs have coevolutionary advantages.

### SC D1 CDR-like loop residues interface with Fcα1-B and the JC

Among the four SC domains, D1 forms the largest interface with JFcα (∼982 Å^2^ buried surface area, not including the SC_N-ext._) using residues in its CDR-like loops (hereafter, CDRs) to contact residues in Fcα1, a linker connecting JC_W1_ and JC_W2_, the JC C-terminus, and at least one tailpiece, Tp_C_ (Fig. 5, A to E). While the D1-Fcα1-B and D1-JC interfaces are mostly conserved when compared to those interfaces in mammalian SIgA, a ∼252 Å^2^ D1-Fcα1-A interface observed in mouse SIgA (also found in human SIgA) is virtually absent in *ap* SJFcα (∼40 Å^2^). Rather than bridging contacts with the Cα2-Cα3 linker and several Cα3 loops, avian D1 CDR2 and surrounding residues are left solvent accessible. This difference appears to stem in part from an orthologous Cα4 loop (residues 373-379) being shorter in most birds (three residues in ducks). Overall, the avian D1 (residues 12-120) makes fewer contacts with JFcα than mammalian D1 makes with IgA; however, the SC_N-ext._ provides an additional ∼506 Å^2^ of buried surface area such that the combined buried surface area of SC_N-ext._ and D1 with JFcα (omitting Tps) is ∼1488 Å^2^ and larger than the mammalian D1-IgA interface (∼1171 Å^2^ in mice, omitting Tps) (fig. S7C).

**Fig. 5.**
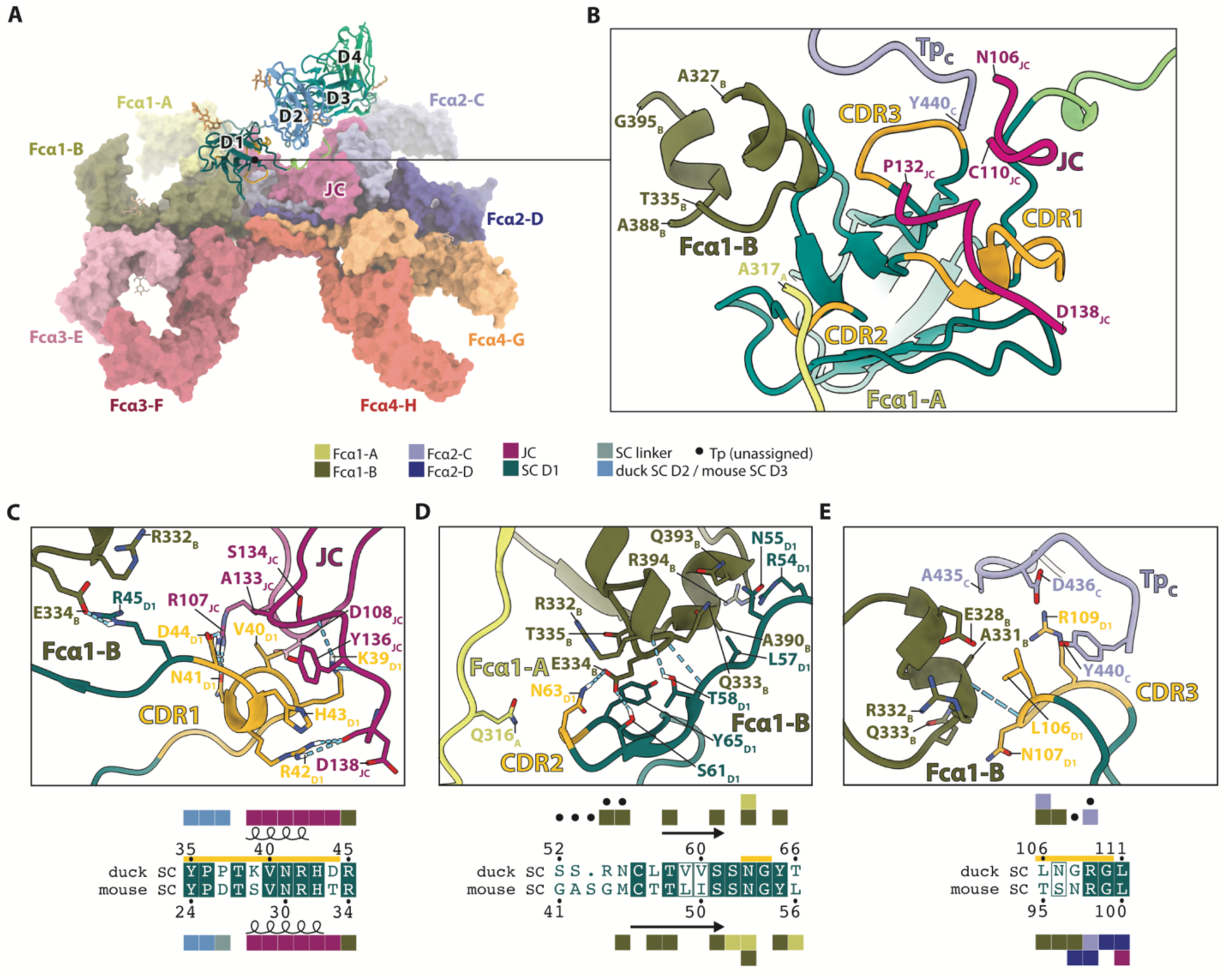
Molecular interfaces of SC D1 in *ap* SJFcα. (**A**) Overall structure of *ap* SJFcα with HCs and JC as surfaces and SC as a cartoon. Components are colored as in Fig. 3. Glycans are shown as sticks. (**B**) Duck SC cartoon showing D1 interfaces with HCs and JC. CDRs are colored golden and numbered. (**C**) Top, cartoon of the duck SC D1 CDR1 interface with Fcα1-B and the JC. Contacting residues are labeled and shown as sticks. Bottom, a structure-based sequence alignment of duck SC D1 CDR1 and mouse SC D1 CDR1. Residue contacts with other chains are indicated by squares colored according to key above. (**D**) Top, duck SC D1 CDR2 interface with Fcα1-A and Fcα1-B. Contacting residues are labeled and shown as sticks. Bottom, a structure-based sequence alignment of duck SC D1 CDR2 and mouse SC D1 CDR2 and surrounding residues with contacts indicated as in (C). Contacts with residues in unassigned Tp chains are indicated by black dots. (**E**) Top, duck SC D1 CDR3 interface with Fcα1-B and Fcα2-C. Contacting residues are labeled and shown as sticks. Bottom, a structure-based sequence alignment of duck SC D1 CDR3 and mouse SC D1 CDR3 with contacts indicated as in (C) and (D).

### D1-D2 linker and D2-D4 anchor SC

Avian pIgR and SC lack a domain homologous to mammalian D2, which protrudes away from D1 and D3; these two domains form a hydrophobic interface (or *clamp*) that holds them together and stabilizes SC as it bridges Fcα1 and Fcα2 (*11, 28*). Despite the absence of D2, we observed D1 and D2 forming an equivalent a hydrophobic interface (∼422 Å^2^) in *ap* SJFcα (Fig. 6, A to C). In contrast to mammals, we observed an additional, smaller (∼132 Å^2^) hydrophobic interface between residues in the first D1 β-strand (A strand; Tyr22, Leu24) and the nine-residue D1-D2 linker (residues 121-129) (Fig. 6, A and C). This smaller interface is separated from the D1-D2 clamp by negatively charged residues in the D1-D2 linker (Glu127, Glu128) and D2 H strand (Glu224, Asp226) and is further stabilized by electrostatic interactions between these residues and D1 Arg19 and D2 Lys223. Overall, the avian D1-D2 interface appears to play a similar role to the D1-D3 interface in mammals, albeit with a more extensive surface area, which may be advantageous for pIgR binding and/or SIgA stability in the absence of a domain homologous to mammalian D2.

**Fig. 6.**
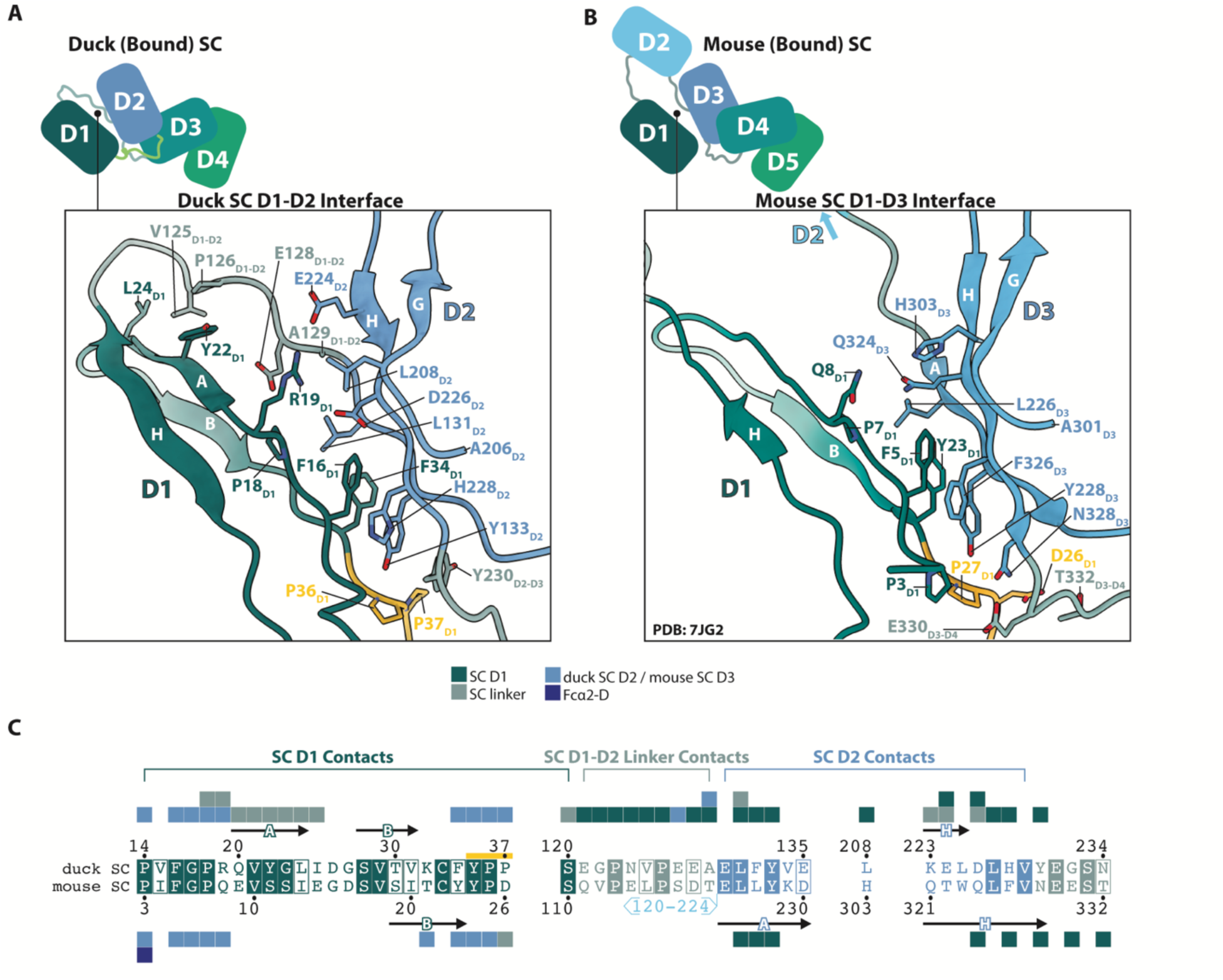
Comparison of duck SC D1-D2 and mouse SC D1-D3 molecular interfaces. (**A**) Schematic of duck SC domain organization in *ap* SJFcα complex with D1-D2 molecular interface shown below as cartoon with contacting residues labeled and shown as sticks. β strands are labeled using Ig nomenclature. (**B**) Schematic of mouse SC domain organization in SIgA complex (PDB: 7JG2) with D1-D3 interface shown below as cartoon with contacting residues labeled and shown as sticks. β strands are also labeled. (**C**) Structure-based sequence alignments comparing duck SC D1-D2 and mouse SC D1-D3 interfacing residues are shown. Residue contacts with other chains are indicated by squares colored according to key.

While D2 shares a significant interface with D1, it makes minimal contact with JFcα. However, residues in D2 (Trp203) and the D2-D3 linker (Glu231) do interface with the SC_N-ext._, thereby forming an interface not found in mammalian SIgA (Fig. 4, A and B). As previously mentioned, the SC_N-ext._ forms its own interface with JC N-terminal residues, including β1 strand residues such as Arg10. We observed JC Arg10 forming a salt bridge with avian D2 Glu204 (Fig. 4B), stabilizing the domain’s position in a location that is shifted ∼3 Å compared to mammalian D3 (measured using the mouse SIgA structure). This difference appears to limit contacts between avian D3, JC_W1_, and the D3-D4 linker compared to mammalian SIgA; mouse D4 residue Glu341 and D4-D5 linker residue Arg443 contact the JC_W1_. The avian D4-JC_W1_ and D4-Fcα2-C interfaces are similar to the corresponding D5-JC_W1_ and D5-Fcα2-C mammalian interfaces, with avian D4 forming additional contacts with Fcα2-C through its DE loop (residues 402-405) (Fig. 7, A to C). Unlike the mouse D5 DE loop, in *ap* SJFcα the D4 DE loop is ordered and possibly stabilized through a glycosylation on Asn404 that is not conserved in mice (Fig. 7, B and C). Overall, the interface formed between SC Ig-like domains and IgA appears to be fairly conserved between mammals and birds with the most apparent difference being the existence of the avian SC_N-ext._ and its resulting contacts.

**Fig. 7.**
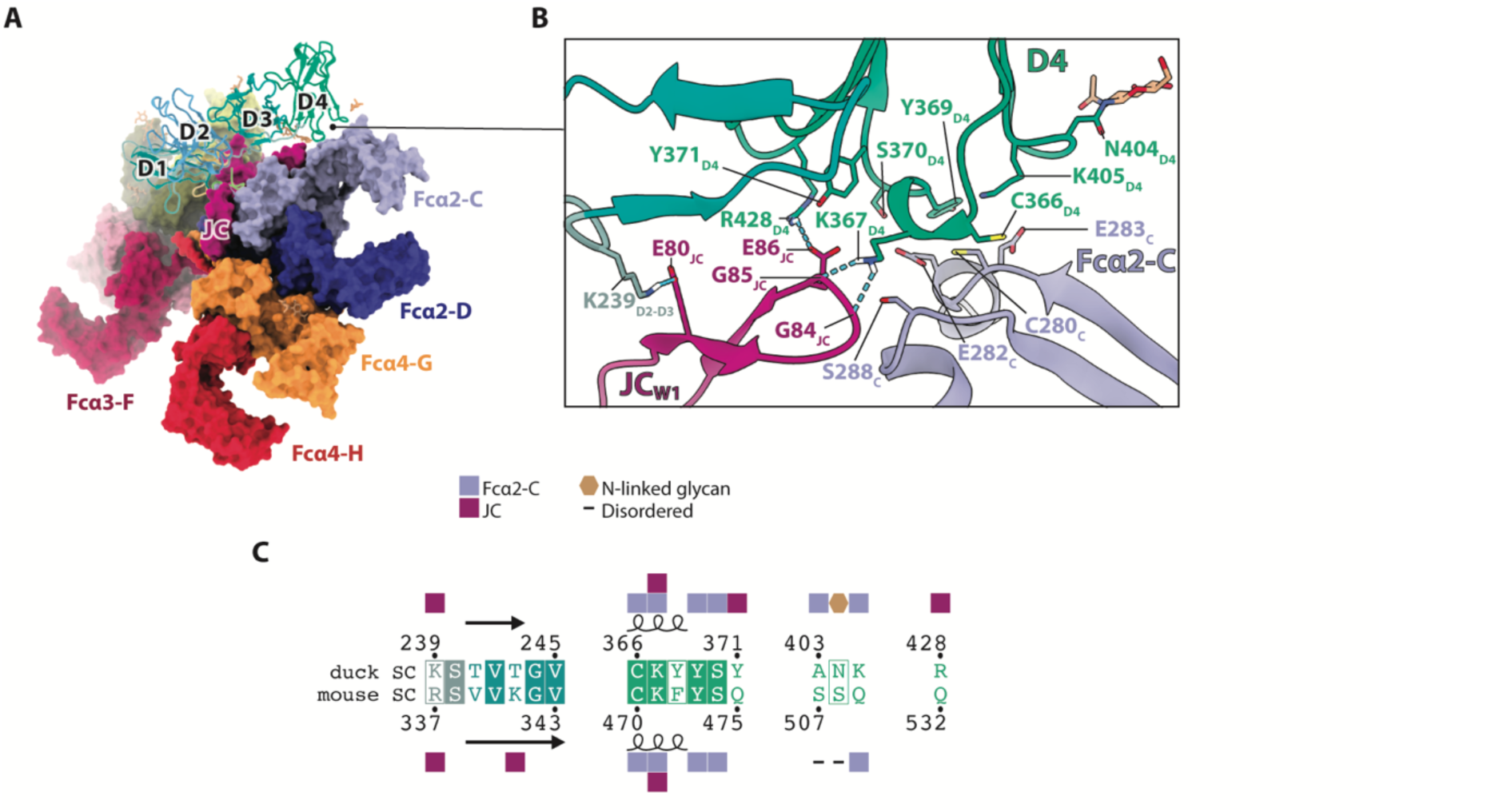
Duck SC D4 molecular interfaces. (**A**) *ap* SJFcα with HCs and JC as surfaces and SC as a cartoon. Components are colored as in Fig. 3. Glycans are shown as sticks. (**B**) Duck SC D4 interface with Fcα2-C and the JC_W1_. Contacting residues are labeled and shown as sticks. N404-linked glycosylation is shown as sticks. (**C**) Structure-based sequence alignment of indicated duck and mouse SC D4 residues. Residue contacts with other chains are indicated by key. Dashes indicate disordered residues.

### Unique avian SC structural features impact IgA binding

Although pIgR binds polymeric IgA in both mammals and birds (*24*, *29*–*31*), the distinct structural features we observed at the SC-JC and SC-JFcα interfaces in *ap* SJFcα prompted us to investigate the molecular mechanisms underlying avian SIgA assembly and compare them to those in mammals using structure-based mutational analysis and surface plasmon resonance (SPR) binding assays. To accomplish this, we expressed recombinant full-length mallard duck (*ap*) SC, mouse (*Mus musculus*; *mm*) SC, and human (*Homo sapiens*; *hs*) SC, as well as variants with modified domain organization, and tested binding to cognate and non-cognate IgA ligands, including *ap* JFcα, *mm* JFcα, and *hs* IgA2. SC and D1 analytes, established soluble models for pIgR-dIgA binding, do not fit single-state kinetic models (*28, 30, 31*); therefore, we qualitatively compared concentration-matched responses and concentration-matched normalized responses for each analyte-ligand pair (Fig. 8; fig. S8). SPR sensorgrams containing all data are shown in figures S9 to S12.

**Fig. 8.**
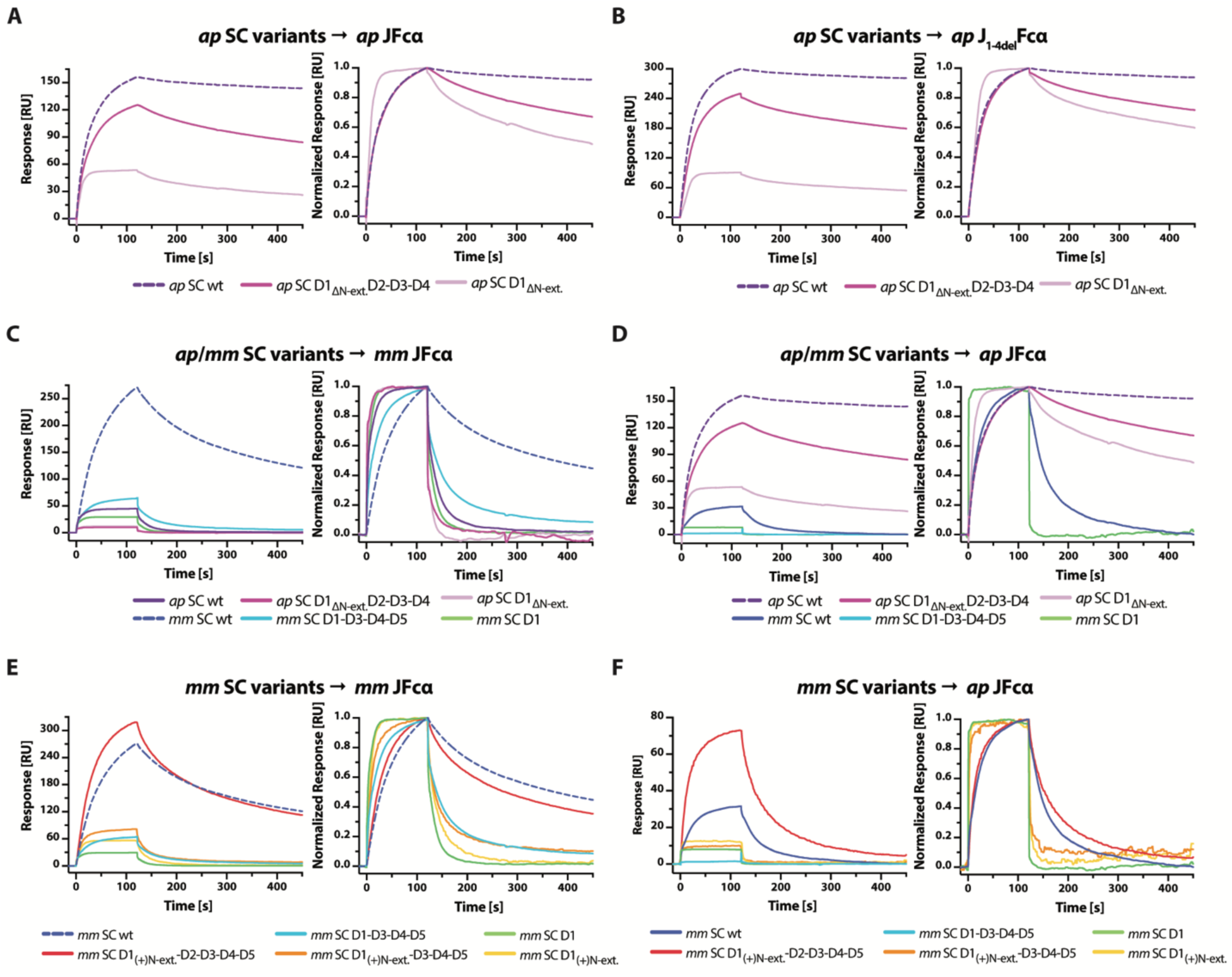
Duck and mouse SC variant binding to cognate and non-cognate ligands. In all panels, SPR sensorgrams showing the response (left) and normalized response (right) of 64nM SC variant analytes binding to JFcα ligands for the following: (**A**) *ap* SC to *ap* JFcα, *ap* SC to *ap* J_1-4del_Fcα, (**C**) *ap* SC and *mm* SC variants to *mm* JFcα, **(D)** *ap* SC and *mm* SC variants to *ap* JFcα, **(E)** *mm* SC and chimeric *mm* SC variants to *mm* JFcα, and (**F**) *mm* SC and chimeric *mm* SC variants to *ap* JFcα. Each analyte response is colored according to the key at the bottom of the panel and dashed lines indicate cognate interactions between unmutated variants. Normalized responses for *mm* SC D1-D3-D4-D5 binding to *ap* JFcα (F) were not calculated due to low responses. Results are consistent with three or more replicate experiments. Sensorgrams for the full analyte concentration series of each experiment are provided in fig. S9-10,12.

To characterize avian SC binding to avian IgA and assay contributions from the SC_N-ext._ and the JC_N-ext._, both of which contribute to unique interfaces, we compared binding of *ap* SC wt and an SC variant lacking the SC_N-ext._ (*ap* SC D1_ΔN-ext._-D2-D3-D4) to immobilized *ap* JFcα and *ap* JFcα containing a JC_N-ext._ deletion (*ap* J_1-4del_Fcα). In SPR experiments with the *ap* JFcα ligand, *ap* SC wt binding responses exhibited moderate association and slow dissociation (Fig. 8A; fig. S9A). In contrast, the *ap* SC D1_ΔN-ext._-D2-D3-D4 sensorgrams exhibited ∼20% lower maximum response units (RU) compared with concentration-matched *ap* SC wt responses (Fig. 8A; fig. S9A), indicating weaker binding. Consistent with this observation, overlaid, concentration-matched, normalized responses revealed a more rapid dissociation phase for *ap* SC D1_ΔN-ext._-D2-D3-D4 compared to *ap* SC wt (Fig. 8A). These results suggest that the SC_N-ext._ is not essential for binding but contributes to avian SC-IgA interactions, possibly stabilizing the *ap* SJFcα complex. To determine if the JC_N-ext._ contributes to SC-IgA binding, we tested binding of *ap* SC wt, *ap* SC D1_ΔN-ext._-D2-D3-D4, and *ap* SC D1_ΔN-ext._ to the *ap* J_1-4del_Fcα ligand. Sensorgram responses and overlaid, concentration-matched, normalized responses for all three analytes were similar to those observed for binding *ap* JFcα ligand (Fig. 8, A and B; fig. S9, A and B), suggesting that the JC_N-ext._ does not significantly impact cognate SC-IgA binding or SIgA complex stability.

We also sought to verify that avian SC D1 is sufficient for binding avian IgA, as has been reported for mammalian and chicken SC variants when binding mammalian IgA (*28–31*). Though we were unable to produce *ap* SC D1 containing the SC_N-ext._, the D1 variant without the SC_N-ext._ (*ap* SC D1_ΔN-ext._) exhibited robust, concentration-dependent binding to *ap* JFcα (Fig. 8A; fig. S9A). Compared to *ap* SC wt and *ap* SC D1_ΔN-ext._-D2-D3-D4, *ap* SC D1_ΔN-ext._ showed more rapid association and dissociation phases (Fig. 8A).

In addition to the SC_N-ext._ and JC_N-ext._ motifs, we identified species-specific contact differences at SC-HC and SC-JC interfaces, despite overall structural conservation. We hypothesized that these differences, along with the absence of an avian SC domain equivalent to mammalian D2, may have evolved to support species-specific, ligand-binding mechanisms. To test this hypothesis, we measured *mm* SC and *ap* SC variants binding to *mm* JFcα and *ap* JFcα ligands and compared the binding profiles from cognate and non-cognate interactions.

We first measured mammalian SC variant binding to cognate ligands. Binding between *mm* SC wt and *mm* JFcα showed moderate association and dissociation whereas *mm* SC D1 and *mm* JFcα binding exhibited comparatively rapid association and dissociation (Fig. 8C; fig. S10A), consistent with previous reports (*28*). We also measured binding between a mouse SC variant lacking D2 (*mm* SC D1-D3-D4-D5) and *mm* JFcα and observed more rapid association and dissociation phases compared to *mm* SC wt (Fig. 8C; fig. S10A). Human variants showed similar trends, where *hs* SC D1-D3-D4-D5 and *hs* SC D1 bound *hs* IgA2 with more rapid association and dissociation phases than *hs* SC wt (fig. S8A; S11A), consistent with prior studies (*28, 30*). For comparison, we measured non-cognate binding of *ap* SC wt, *ap* SC D1_ΔN-ext._-D2-D3-D4, and *ap* SC D1_ΔN-ext._ to *mm* JFcα, which revealed rapid association and dissociation compared to equivalent experiments testing binding to *ap* JFcα (Fig. 8C; fig. S10B). A similar pattern was observed for *ap* SC wt and *ap* SC D1_ΔN-ext._ binding to *hs* IgA2 (fig. S8A; S10E) though differences between *ap* SC wt and *hs* SC wt binding to *hs* IgA2 were less pronounced and reflected more moderate changes to the dissociation phase. Collectively, results reinforce a role for D2 in supporting cognate mammalian SC-IgA interactions and indicate that differences between duck, mouse, and human IgA ligands can impact avian SC binding.

Next, we tested non-cognate interactions between mammalian SC variants and *ap* JFcα. Sensorgrams associated with *mm* SC wt binding to *ap* JFcα exhibited lower overall responses and markedly more rapid dissociation phases compared to cognate *ap* SC wt binding to *ap* JFcα (Fig. 8D; fig. S12A). SPR profiles were similar for *hs* SC wt binding to *ap* JFcα (fig. S8B and fig. S12B). We observed low responses for *mm* SC D1-D3-D4-D5 and *mm* SC D1 binding to *ap* JFcα (∼1 RU and ∼7 RU, respectively), indicating relatively weak interactions (Fig. 8D; fig. S12A). SPR responses for *hs* SC D1-D3-D4-D5 and *hs* SC D1 binding to *ap* JFcα were also reduced (∼34 RU and ∼29 RU, respectively), and displayed rapid association and dissociation compared to *mm* SC wt and *ap* SC wt (fig. S8B and S12B). Notably, the binding profiles of *mm* SC D1-D3-D4-D5 and *hs* SC D1-D3-D4-D5 to *ap* JFcα did not resemble that of *ap* SC wt, indicating that deletion of D2 was insufficient to reproduce avian SC binding profiles. The markedly low responses associated with *mm* SC D1-D3-D4-D5 and *mm* SC D1 binding to *ap* JFcα were unexpected because our data demonstrate that *mm* SC D1-D3-D4-D5 can bind *mm* JFcα (Fig. 8C; fig. S10A) and a majority of the D1 CDR contacts with IgA are conserved between mammals and birds (Fig. 5, C to E). However, contacts between SC and IgA outside of D1 and/or conformational differences between IgA ligands may have a greater influence on non-cognate binding. Together, our observations suggest species-specific differences in SC-IgA ligand recognition and SIgA complex stability.

Results highlighting differences in SC variant binding to cognate and non-cognate ligands, along with the observation that mammalian D2 deletion did not recapitulate avian SC binding profiles, suggested more complex coevolutionary relationships between SC and IgA. To test whether transferring additional avian SC features to mammalian SC variants could recapitulate *ap* SC wt binding to *ap* JFcα, we engineered chimeric mammalian SC constructs, in which the mallard duck SC_N-ext._ was fused to the N-terminus (*mm* SC D1_(+)N-ext._-D2-D3-D4-D5, *mm* SC D1_(+)N-ext._-D3-D4-D5, *mm* SC D1_(+)N-ext._, *hs* SC D1_(+)N-ext._-D2-D3-D4-D5, *hs* SC D1_(+)N-ext._-D3-D4-D5, and *hs* SC D1_(+)N-ext._) and measured binding to the corresponding mammalian ligand and *ap* JFcα. SPR sensorgrams for chimeric mouse SC variants binding to *mm* JFcα resembled those of non-chimeric variants; however maximum responses at 64 nM were 17%, 25%, and 93% higher, respectively, indicating that the SC_N-ext._ can enhance mouse SC binding to binding to mm JFcα (Fig. 8E; fig. S10C). In contrast, sensorgrams of chimeric human SC constructs binding to *hs* IgA2 were comparable to non-chimeric variants (fig. S8C and S11C). These results suggest that while the SC_N-ext._ improves mouse SC binding to mouse JFcα, it has less impact on human SC-IgA2 interactions. We also tested chimeric mouse SC variant binding to *ap* JFcα, which exhibited higher maximum RU than non-chimeric variants, and for *mm* SC D1_(+)N-ext._-D2-D3-D4-D5, moderately slower dissociation compared to non-chimeric variants (Fig. 8F; fig. S12C). Notably, *mm* SC D1_(+)N-ext._-D3-D4-D5 *and mm* SC D1_(+)N-ext._ exhibited concentration dependent binding to *ap* JFcα, whereas equivalent variants lacking the SC_N-ext._ exhibited very weak responses (Fig. 8F; fig. S12C). Experiments using chimeric human SC variants produced similar trends, showing enhanced binding to *ap* JFcα compared to non-chimeric equivalents (fig. S8F and S12D). Together, results suggest adding the avian SC_N-ext._ to mammalian SC can enhance binding to the non-cognate *ap* JFcα ligand.

## DISCUSSION

IgA evolved to become the predominant mucosal antibody in modern birds and mammals, and its structures are broadly relevant to understanding antibody evolution and elucidating how structural differences among antibody classes and species correlate with unique protective functions. Across species, IgA shares the ability to form JC-containing multimers with IgM (except for teleosts which have lost the JC). In birds, IgA and IgM also share HC domain organization, with IgA containing four constant domains (Cα1-Cα4), including a domain equivalent to Cμ2. This latter avian IgA feature distinguishes it from mammalian IgA, which lacks a HC domain equivalent to avian Cα2. Polymeric state varies among JC-containing antibodies; while JC-containing IgM is typically pentameric, IgA is predominantly dimeric in mammals, with additional variability across species and isoforms. Whereas endogenously and recombinantly expressed human pIgA contains up to 10% tetrameric IgA (*1, 20*), we found that 90% or more of recombinantly expressed *ap* JFcα is tetrameric, consistent with earlier studies indicating that endogenous mallard duck IgA is tetrameric (*23*). Together, these observations raise the question of how the avian tetrameric IgA, with its additional HC domain, functionally differs from mammalian IgM and IgA.

Our structures reveal *ap* JFcα to be a relatively asymmetric structure, with varying degrees of twist and tilt between JC-linked monomers. The IgA asymmetry may be “fixed,” as observed in the structure, or could in part arise dynamically, where “breathing” at the interfaces results in conformational variability. This asymmetry differs from human IgM structures, that reveal relatively flat, or planar, pentameric Fcμ cores. Together, avian and mammalian IgA structures broadly suggest that the conformations of polymeric IgAs are typically less planar than IgM. Varying degrees of twist and tilt between JC-linked monomers are observed in human IgA structures, including the low-abundance human SIgA1 tetramer (*10*). Our data indicate that avian Cα2, which is absent in mammalian IgA, may further increase the asymmetry of the avian complex. Specifically, our data demonstrate that positions occupied by avian Cα2 extend the twist and tilt observed at the core of the molecule toward the Fabs, likely enforcing a distinct geometric relationship between the four Fabs (and their antigens) compared to mammalian tetrameric IgA. The presence of avian Cα2 may also allow for Fab flexibility in avian IgA that is distinct from human IgA. Structural studies of human IgM indicate that Cμ2 domains are flexible, albeit to varying extents, allowing the Fabs to move relative to the IgM core (*32*). Avian IgA’s tetrameric conformation, combined with potential Cμ2-Fab conformational flexibility may promote high avidity antigen interactions and/or antigen crosslinking. Antigen cross-linking mechanisms (e.g. classical agglutination or enchained growth) are especially critical for SIgA-dependent immune exclusion, a process in which mucosal antibodies keep pathogens away from epithelial barriers and promote removal via physical processes such as peristalsis. We speculate that avian IgA’s predominantly tetrameric state may enhance these mechanisms compared to dimeric mammalian IgA and may represent an evolutionary adaptation to the distinct pathogens encountered by birds. Consistent with this idea, human SIgA tetramers have been reported to have greater activity against some viruses compared to SIgA dimers, although the underlying mechanisms behind protection are not well understood (*33–35*).

The pIgR is central to mucosal antibody function, promoting transcytosis and release of JC-containing SIgM and SIgA into the mucosa (*14*). This process is conserved in birds and mammals, and in the case of SIgM, extends to amphibians and some reptiles (*6*, *24*, *3C*). However, as noted, pIgR domain organization differs among species (*C*) and our *ap* SJFcα structure revealed that these differences resulted in structural variations between liganded SC in birds and mammals. Perhaps the most notable differences are the SC and JC N-terminal extensions. Our sequence alignments show that an SC_N-ext._ is found in all representative bird species (fig. S7A), although conservation is variable. An SC_N-ext._ also appears to exist in turtles; however, the length and sequence is not conserved. Furthermore, our alignments indicate that a JC_N-ext._ is found in many non-mammalian species as well as the platypus, but the length and residue identity are variable (fig. S7B). Although we found the JC_N-ext._ is stabilized when SC is present (disordered in *ap* JFcα), thereby suggesting a structural role in SIgA, these extensions may have other functionals roles (e.g. binding to other receptors when SC is absent). Regardless, their presence in birds suggests they provide an advantage associated with avian IgA and/or IgM functions. Consistent with this, our *ap* SJFcα structure indicates that these extensions increase the shared surface area between SIgA components and, in the case of SC, influence binding kinetics. While the SC and JC N-terminal extensions were not required for binding, deletion of the SC_N-ext._ appeared to impact dissociation, suggesting a role in maintaining complex stability. This is further supported by our results demonstrating that adding the SC_N-ext._ to mammalian SC variants promoted binding. Although a role in IgA binding and stability is apparent for the SC_N-ext._, its contribution to broader *in vivo* functions remains unknown, but it could include roles in avian SIgM assembly and/or stabilization of unliganded pIgR. Moreover, while we observe a clear structural role for the JC_N-ext._, its deletion did not markedly impact avian SC binding, reinforcing the possibility that it functions in SIgM assembly, promotes complex stability in conditions not assayed in our experiments (e.g. mucosal secretions and timeframes exceeding 400 seconds), and/or is involved in binding to other factors (e.g. FcRs).

Avian SC is known to lack a domain homologous to mammalian D2 (*31, 37*). In mammalian SIgA structures, SC D2 extends away from the complex while the other SC domains form contacts with IgA HCs and JC. Here we find that avian SC forms contacts with Fcα1, Fcα2, and JC that are largely equivalent to those observed in mammalian structures (excluding noted contributions from SC_N-ext._), indicating that the absence of a domain equivalent to mammalian D2 does not markedly alter the SC-IgA interactions in birds. It is also notable that deleting D2 from mammalian SC sequences markedly impacts their interactions with IgA and IgM (*30*) (Fig. 8C; fig. S8A), resulting in apparently weaker binding than avian SC, which naturally lacks a domain equivalent to D2. Notably, adding the SC_N-ext._ to mammalian SC variants lacking D2 did not result in binding profiles comparable to *ap* SC wt. Collectively, these results indicate that species-specific differences, though not yet experimentally tested, impact cognate SC-IgA binding kinetics.

Whereas the role of pIgR in transporting IgM and IgA is well established, the role of liganded SC in mucosal secretions is less clear (*1, 11, 14*). The *ap* JFcα and *ap* SJFcα structures revealed subtle differences in the JFcα conformation when SC is bound, characterized by more negative tilt values. While it remains unknown if these changes are functionally significant, in principle, they may impact Fab interactions with antigen and/or alter the accessibility of immune cell receptor binding sites. We also note that with only four domains, avian SC has a reduced solvent-accessible surface area and fewer glycosylation sites (four) when compared to mammalian SC (seven), features which have been proposed to mediate SC functions in mammals (*1*).

The conformations of the *ap* JFcα and *ap* SJFcα structures are also likely to impact putative avian IgA interactions with immune effector cells. In humans, IgA binding to FcαRs, including FcαR1 (CD89) and FcαμR, can initiate IgA-dependent effector functions. Recent structural studies suggest that the ability of CD89 to interact with SIgA-antigen complexes is dependent on the polymeric state of IgA (*1*, *16*); less is known about human FcαμR structure and function. Avian FcαRs remain uncharacterized and thus it is unclear if and how they would interact with avian IgA, although our analysis indicates that the IgA residues forming the human CD89 binding site are not conserved in *ap* JFcα (not shown). Moreover, the tetrameric structure may limit steric accessibility to those sites compared to monomeric and dimeric forms of IgA found in mammals. The mammalian IgM FcR, FcμR, can bind both faces of IgM depending on the presence of SC. While in principle an avian IgA receptor could bind in a similar manner, we also find that a majority of the human IgM residues involved in human FcμR binding are not conserved in *ap* JFcα (not shown). These observations lead us to speculate that putative avian FcαRs target distinct bindings sites compared to mammalian variants, raising the possibility that cell-based effector mechanisms differ markedly in birds.

Understanding how different species’ immune systems counteract similar pathogens is critical for combatting zoonotic disease. This comparison is especially relevant for humans and aquatic birds, including many species of ducks, which are frequently exposed to avian influenza virus yet rarely display symptoms when infected with strains that cause severe symptoms in humans (*38*) and other avian species, like chickens (*39*–*41*). Because influenza enters through a host’s mucosal surfaces (i.e. human upper respiratory tract; duck gastrointestinal tract) where SIgA serves as the front-line antibody defense (*23, 42*), we speculate that the structural differences we observed provide ducks with an advantage. Specifically, we envision that avian SIgA provides an asymmetric, tetrameric platform, unique to birds, that when combined with the unique duck antibody repertoire could provide exceptional protection against pathogens. This is supported by the observation that human tetrameric IgAs have been associated with increased efficacy against flu (*33–35*) and the observation that duck immune systems and influenza viruses are thought to have a strong co-evolutionary history (*7*).

## MATERIALS AND METHODS

### Study design

The objective of this study was to structurally characterize secretory immunoglobulin A in birds. Cryo-EM was used to determine the structures of *ap* JFcα and *ap* SJFcα at 3.79-Å and 3.21-Å resolutions, respectively. SPR was used to characterize SC-IgA interactions and the relative contributions from structural motifs identified in this study.

### Construct design

Protein sequences for the *Anas platyrhynchos* (*ap*; mallard duck) IgA heavy chain (HC) constant region (GenBank: AAA68606), joining chain (JC, NCBI: XP_005031370), and pIgR (NCBI: XP_021122629), as well as for *Mus musculus* (*mm*; house mouse) IgA HC constant region (UniProt: P01878), JC (UniProt: P01592), and pIgR (UniProt: O70570), and *Homo sapiens* (*hs*; human) IgA2 HC constant region (NCBI: AAT74071.1), light chain constant region (LC, UniPROT: P0CG04), JC (NCBI: NP_653247.1), and pIgR (UniProt: P01833) were obtained from the NCBI database. The sequences encoding the *Anas platyrhynchos* IgY (PIR: B465529) signal peptide (MSPRPHAFALLLLLAAVPGLRA) (*43*), a N-terminal hexa-histidine tag, and mallard duck IgA HC residues 106 to 440 (Cα2-Cα3-Cα4-Tp) were fused, codon optimized for human cell expression, synthesized (Integrated DNA Technologies, Inc.), and cloned into mammalian expression vector pD2610-v1 (Atum). The mouse IgA HC construct was generated as previously described (*28*).

The human IgA2 HC was fused to the tPA signal peptide and the STA121 HC variable region (*44*), and cloned into pD2610-v13 (Atum). The human LC construct was cloned into pD2610-v1 and included the mouse Ig κ signal peptide (METDTLLLWVLLLWVPGSTG), the STA121 LC variable region (*44*), and the human LC constant residues fused together.

JC and pIgR ectodomain (secretory component, SC) constructs retained their endogenous signal peptides. Sequences encoding mallard duck JC (residues 1 to 138), mallard duck JC with JC_N-ext._ deletion (residues 5 to 138), mouse JC (residues 1 to 137), mallard duck SC (residues 1-458 and a C-terminal hexa-histidine tag), and mouse SC (residues 1 to 567 and a C-terminal hexa-histidine tag) were codon optimized, synthesized, and cloned into pD2610-v1.

For surface plasmon resonance (SPR) experiments, mallard duck SC (*ap* SC wt), mouse SC (*mm* SC wt), and human SC (*hs* SC wt) sequences were modified and synthesized to create the following constructs in pD2610-v1 (Atum): mallard duck SC without the SC_N-ext._ (*ap* SC D1_ΔN-ext._-D2-D3-D4), mallard duck SC D1 only without the SC_N-ext._ (*ap* SC D1_ΔN-ext._), mouse SC with D2 deletion (*mm* SC D1-D3-D4-D5), mouse SC D1 only (*mm* SC D1), human SC with D2 deletion (*hs* SC D1-D3-D4-D5), and human SC D1 only (*hs* SC D1). Chimeric SC constructs were designed by fusing the mallard duck SC endogenous signal peptide and SC_N-ext._ sequences to the mouse and human sequences listed above and synthesized to create the following constructs in pD2610-v1 (Atum): *mm* SC D1_(+)N-ext._-D2-D3-D4-D5, *mm* SC D1_(+)N-ext._-D3-D4-D5, *mm* SC D1_(+)N-ext._, *hs* SC D1_(+)N-ext._-D2-D3-D4-D5, *hs* SC D1_(+)N-ext._-D3-D4-D5, and *hs* SC D1_(+)N-ext._. To prevent covalent analyte-ligand interactions, C366A/C400A (mallard duck), C470A/C504A (mouse), and C468A/C502A (human) mutations were introduced into all SC constructs used for SPR. DNA and amino acid sequences for all constructs are provided in table S2.

### Protein expression and purification

DNA constructs were transiently transfected into HEK Expi293 cells (Gibco: A14527) using the ExpiFectamine 293 Transfection Kit (Gibco: A14525). Six days after transfection, supernatants were harvested; *ap* JFcα (mallard duck IgA HC and JC),*ap* SJFcα (mallard duck IgA HC, JC, and SC), *mm* JFcα (mouse IgA HC and JC), and SC proteins were purified by affinity chromatography using Ni-NTA Agarose (Qiagen) and eluted with elution buffer containing 50 mM Tris-HCl, 150 mM NaCl, pH 7.4, and 250 mM imidazole. *hs* IgA2 (human IgA2 HC, LC, and JC) was purified using CaptureSelect^TM^ Human IgA Affinity Matrix (Thermo Fisher) and eluted with 0.1 M glycine, pH 3.0. All proteins were further purified using size-exclusion chromatography (Superose 6 Increase 10/300, Cytiva) and elution fractions containing protein were collected and stored in 50mM Tris-HCl, 150 mM NaCl, pH 7.4 at 4°C.

#### Cryo-EM grid preparation and data collection

*ap* JFcα: UltrAuFoil R1.2/1.3 300 mesh grids (Quantifoil) were glow discharged in a PELCO easiGlow (Ted Pella Inc.) for 90 s at 25 mA current. Using a Vitrobot Mark IV (Thermo Fisher Scientific), three applications of 3 uL of sample at 0.1 mg/mL were applied to each grid at 4°C and 100% relative humidity. Each application was followed by a 3 s wait and 2 s blot time. After the third blot, the grid was vitrified. Movies were collected at Purdue University on a Titan Krios G4 (Thermo Fisher Scientific) operating at 300 kV and equipped with a post-GIF K3 direct electron detector (Gatan). 2,544 movies were collected with an untilted stage using EPU (Thermo Fisher Scientific) and 3,015 movies were collected with a stage tilted at 30° using Leginon (*45*). Collections were performed at 81,000x magnification in super resolution mode with total exposure time of 3.21 s, total dose of 57.79 electrons/Å^2^, raw pixel size of 0.527 Å/pixel, and 40 frames per movie.

*ap* SJFcα: Holey carbon R1.2/1.3 400 mesh copper grids (Quantifoil) were glow discharged for 60 s at 25 mA current in a PELCO easiGlow (Ted Pella Inc.). Using a Vitrobot Mark IV (Thermo Fisher Scientific), 1 blot of 3 uL of sample at 0.4 mg/mL was applied to each grid at 4°C and 100% relative humidity with 10 s wait time, 2 s blot time, and 0 s drain time. Movies were collected on a Titan Krios G4 (Thermo Fisher Scientific) at Purdue University operating at 300 kV and equipped with a post-GIF K3 direct electron detector (Gatan). 3,009 movies were collected with an untilted stage using EPU (Thermo Fisher Scientific) and 4,034 movies were collected with a stage tilted at 30° using SerialEM (*46*). Collections were performed at 105,000x magnification with total exposure time of 2.41 s, total dose of 71.3 electrons/Å^2^, raw pixel size of 0.822 Å/pixel, and 60 frames per movie.

#### Cryo-EM data processing

All data processing was performed in cryoSPARC (*47*).

*ap* JFcα: Raw movie frames were motion corrected with an output F-crop factor of 1/2 and contrast transfer function (CTF) corrected. Micrographs were screened based on CTF fit, ice thickness, and presence of contamination using the Manually Curate Exposures module to remove poor-quality micrographs, resulting in 2,180 untilted and 2,915 tilted micrographs. The Blob picker module was used to pick particles with a minimum and maximum particle diameter of 150 Å and 300 Å from approximately half of the untilted micrographs. Picked particles were subjected to 2D classification to generate templates for template-based particle picking. The Template picker module was used to pick particles from untilted and stage-tilted datasets with a diameter of 230 Å and minimum particle separation of 0.2 diameters. The 10.6 M picked particles underwent several rounds of 2D classification, resulting in 614,430 particles, which were used for initial model generation and refinement using Ab-Initio Reconstruction and Non-uniform Refinement, respectively. The final refinement generated a map using 230,667 particles with an overall resolution of 3.76 Å (Fourier shell correlation (FSC) = 0.143).

*ap S*JFcα: Raw movie frames were motion and CTF corrected. Micrographs were screened as described above, resulting in 2,958 untilted and 3,777 tilted micrographs. Blob picker module was used to pick particles with a minimum and maximum particle diameter of 200 Å and 400 Å from untilted and tilted micrographs. Picked particles from untilted and tilted groups underwent separate rounds of 2D Classification. Resulting 2D classes from the two groups were combined and used for Template-based particle picking. Picked particles were processed as described above. The final refinement generated a map using 688,138 particles with an overall resolution of 3.21 Å (FSC = 0.143).

### Model building, refinement, and validation

trRosetta (*48*) was used to generate a de novo model of mallard duck HC Cα3 and Cα4 domains. Individual domains were docked into the cryo-EM maps using UCSF ChimeraX 1.8 (*4S*). Initial inspection of the docked model revealed most loop regions fit poorly into the density; these regions were deleted from the model and then manually rebuilt using Coot 0.9.8.1 (*50*). IgA tailpieces, the JC (residues 8-138), and SC D1-D4 were manually built using Coot while referencing previously published polymeric antibody structures (*10, 11*). The mallard duck JC N-terminus (residues 1 to 7) and SC_N-ext._ (residues 1 to 11) were built de novo using Coot. Real space refinement was performed using Coot and Phenix (*51*).

### Structural analysis

To quantify the geometric relationships between the four Fcα subunits, the *ap* JFcα structure was imported into UCSF ChimeraX and the built-in “Axes/Planes/Centroids” tool was used to define a fixed (JFcα) coordinate system (fig. S4B) and a local (Fcα subunit) coordinate system (fig. S4C). Axes, planes, and centroids were defined using residues located in the eight HC Cα3 and Cα4 domains (residues 213 to 418) for the fixed coordinate system and using residues located in the intra-subunit pairs of Cα3 and Cα4 domains (residues 213 to 418) for the local coordinate systems. Origins for each coordinate system were defined as the non-mass weighted centroid for selected residues. The x, y, and z axes of each coordinate system were defined as a centroid-anchored axis that aligned with either the primary, tertiary, or secondary eigenvector of the selected residues, respectively. Planes were also centroid anchored and aligned with the primary and secondary eigenvectors.

The angle between adjacent Fcα subunits was measured in UCSF ChimeraX using the “Angles/Torsions” tool and defined as the central angle formed between adjacent Fcα subunit centroids and the JFcα centroid (fig. S4D).

To measure the twist and tilt of each Fcα subunit, the local (Fcα subunit) coordinate systems were aligned with the fixed (JFcα) coordinate system (fig. S4E). First, the local (Fcα subunit) coordinate system was rotated about the fixed y-axis (y^JFcα^) until the x-coordinate of the Fcα subunit centroid in the fixed coordinate system was 0. The local coordinate system was then translated to align the local origin (Fcα subunit centroid) with the fixed origin (JFcα centroid). Twist was defined as the degrees (α) of rotation about the fixed z-axis (z^JFcα^) of the local coordinate system (x^Fcα^, y^Fcα^, z^Fcα^) required to form a 90° angle between the local y-axis (y^Fcα^) and the fixed x-axis (x^JFcα^) (fig. S2F). Tilt was defined as the degrees (β) of rotation about the fixed x-axis (x^JFcα^) after the twist rotation of the local coordinate system (x’^Fcα^, y’^Fcα^, z’^Fcα^) required to form a 90° angle between the local y-axis (y’^Fcα^) and the fixed z-axis (z^JFcα^) (fig. S2F).

To compare the spacing between adjacent Fcα subunits, the UCSF ChimeraX “distance” command was used to measure the distance between the carbonyl carbon of residue Cys400 in adjacent Cα4 domains. Buried surface areas were analyzed using PISA (*52*) and UCSF ChimeraX (*49*). Interchain contacts at molecular interfaces were analyzed using PISA (*52*), UCSF ChimeraX (*49*), and ENDscript (*53, 54*).

The “super” command in Pymol (*55*) was used to calculate RMSD between JC-Tailpiece assembly Cα atoms in *ap* JFcα (HC residues 425 to 433, JC residues 8 to 68) and either mouse dIgA (PDB 7JG1; HC residues 452 to 460, JC residues 3 to 63) or human SIgA (6UE8; HC residues 457 to 465, JC residues 4 to 63) structures. This strategy was also used to calculate RMSD between Cα atoms in *ap* JFcα and *ap* SJFcα (with SC removed).

The β strands in each Ig-like domain of SC were assigned a letter consistent with Ig-fold nomenclature (*56*).

### Sequence alignments

pIgR/SC and JC sequences (respectively) were obtained for the following species from NCBI (unless otherwise noted): mallard duck (*Anas platyrhynchos*; XP_021122629; XP_005031370), chicken (*Gallus gallus*; NP_001038109.1; NP_989594.1), emu (*Dromaius novaehollandiae*; XP_025967825.2; XP_025959286.2), malkoha (*Phaenicophaeus curvirostris*; XP_025967825.2; XP_069710933.1), gull (*Chroicocephalus ridibundus*; XP_063212801.1; XP_063191950.1), crow (*Corvus brachyrhynchos*; XP_017596648.1; XP_008635627.1), frog (*Nanorana parkeri*; XP_017596648.1; XP_018409072.1), toad (*Spea bombifrons*; XP_053311900.1; XP_053317628.1), turtle (*Pelodiscus sinensis*; XP_025034348.1; ACH57092.1 (GenBank)), platypus (*Ornithorhynchus anatinus*; XP_007668461.1; XP_003430954.1), mouse (*Mus musculus*; O70570 (Uniprot); P01592 (UniProt)), and humans (*Homo sapiens*; P01833 (UniProt); NP_653247.1). Mature protein sequences were identified using SignalP 6.0 (*57*) before performing sequence alignments using ClustalOmega (*58*). ENDscript (*53*, *54*, *59*) was used to map contacts of JC and/or SC with other chains and assign secondary structure for *ap* SJFcα and mouse SIgA (PDB: 7JG2). The alignment figure panels were generated with ENDscript (*53*, *54*, *59*) and further formatted in Adobe Illustrator.

### Surface plasmon resonance

Surface plasmon resonance (SPR) binding studies were performed using a Sierra SPR-32 Pro (Bruker) operating at 25°C. Ligands were diluted in sodium acetate buffer of either pH 4.0 (*ap* JFcα, *ap* J_1-4del_Fcα, and *mm* JFcα) or pH 4.5 (*hs* IgA2) and immobilized on a High Capacity Amine Sensor (Bruker) using an amine coupling kit (Bruker). Each ligand was immobilized in a separate flow channel with at least one mock-coupled spot per channel to serve as a reference surface.

A two-fold dilution series of each analyte was prepared using HBS-EP+ buffer (10mM HEPES, 150 mM NaCl, 3mM EDTA, 0.05% (v/v) Tween 20) starting at 512 nM, except for *ap* SC D1_ΔN-ext._ with *ap* J_1-4del_Fcα ligand, which started at 64 nM. Analytes were injected in HBS-EP+ buffer at a flow rate of 50 μL/min, with a 120 s association phase and a 320 s dissociation phase. Each injection was followed by a regeneration step (2.5 M MgCl_2_, flow rate of 25 μL/min, 60 s contact time) and two wash steps (HBS-EP+, 60 s contact time each).

Sensorgrams were reference-surface and buffer subtracted in Bruker Sierra Analyzer 3 software and then exported and replotted in GraphPad Prism 10.5.0 for macOS. Normalization of the respective sensorgrams for concentration-matched samples were carried out in Microsoft Excel by dividing RU by the maximum measured response of the respective analyte and then replotted in GraphPad Prism 10.5.0 for macOS. Sensorgrams shown in figures are representative of replicate experiments.

### Figures

Figures were prepared using UCSF ChimeraX and Adobe Illustrator.

## Acknowledgements

The cryo-EM data collection was performed at the Purdue Cryo-EM Facility with the assistance of Thomas Klose and Frank Vago. Asta Simonovic and Sarah Leonard assisted with protein expression and purification. Authors thank Profs. Wilfred van der Donk, Nicholas Wu, and Jenna Guthmiller and all members of the Stadtmueller Lab for insightful discussions; we also thank Sarah Leonard for assistance with SPR instrumentation.

## Author contributions

The study was conceived by BMS and RMS; experiments were conducted by RMS; QL assisted with cryo-EM data collection and processing; and all authors contributed to data analysis and manuscript writing.

## Data and Materials availability

Cryo-EM density maps have been deposited in the EM databank (www.ebi.ac.uk/emdb) with the accession codes EMD-47901 and EMD-47900 for *ap* JFcα and *ap* SJFcα, respectively. The refined coordinate has been deposited in the Protein Data Bank (www.rcsb.org) with accession codes 9EC7 and 9EC6 for *ap* JFcα and *ap* SJFcα, respectively.

## Supplementary Materials

**Fig. S1.**
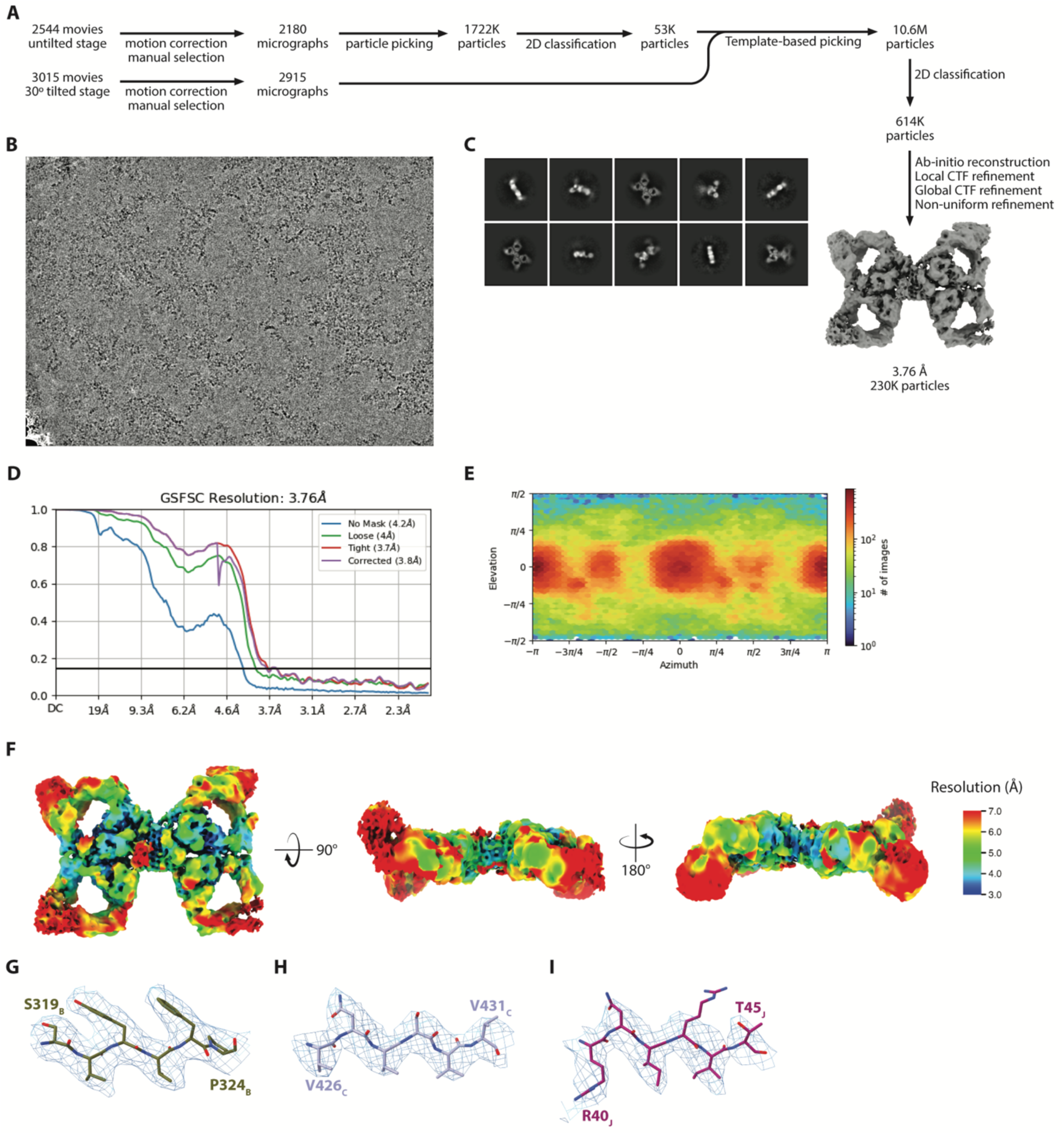
Cryo-EM data collection and CryoSPARC processing pipeline for *ap* JFcα. (**A**) Schematic summary of the *ap* JFcα Cryo-EM data processing pipeline in CryoSPARC. (**B**) Representative micrograph of *ap* JFcα. (**C**) Representative 2D class averages. (**D**) The FSC curve for the final reconstruction with reported resolution at FSC=0.143 shown by the black horizontal line. (**E**) Viewing direction distribution plot of particles used in final refinement. (**F**) Local resolution map calculated by CryoSPARC and colored in ChimeraX. (**G-J**) Cryo-EM map density surrounding residues 319 to 324 in Fcα1-A Cα4 (G), residues 426 to 431 in Tp_C_ (H), and residues 40 to 45 in JC_core_ (I) is contoured to threshold of 0.17 and carved at 2 Å.

**Fig. S2.**
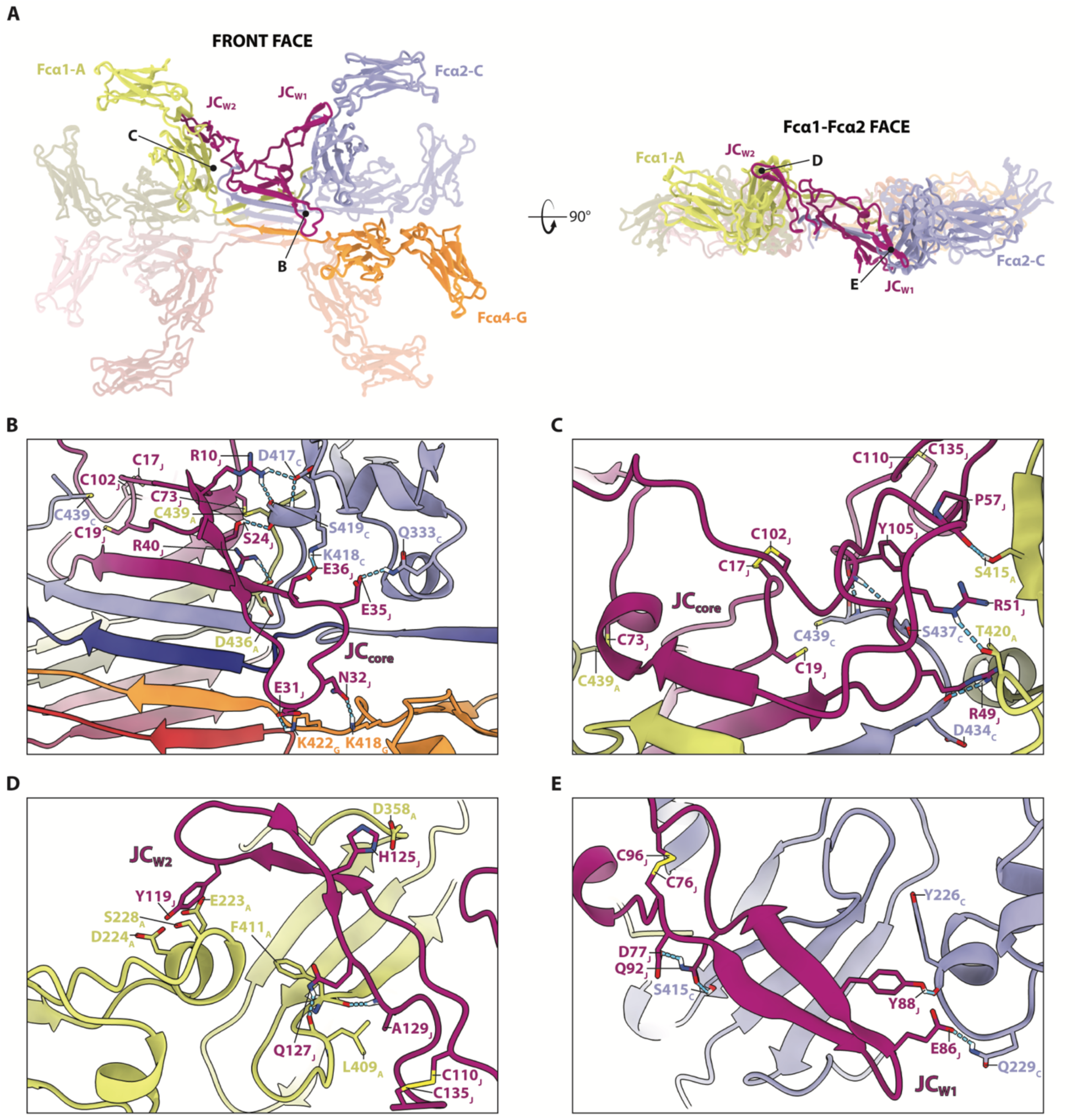
Molecular interactions at *ap* JFcα interfaces between JC and HCs. (**A**) Front face (left) and Fcα1-Fcα2 face (right) views of *ap* JFcα shown as cartoon representation colored as in (1A) with interacting chains labeled. Black lines and letters indicate focused regions for subsequent panels. (**B-C**) Cartoon representation of front face (B) and back face (C) views of polar and electrostatic interactions between JC_core_ and HCs. (**D**) Cartoon representation showing polar and electrostatic interactions between JC_W2_ and Fcα1-A. (**E**) Cartoon representation showing polar and electrostatic interactions between JC_W1_ and Fcα2-C. Interacting residues and relevant Cys residues are labeled and shown as sticks.

**Fig. S3.**
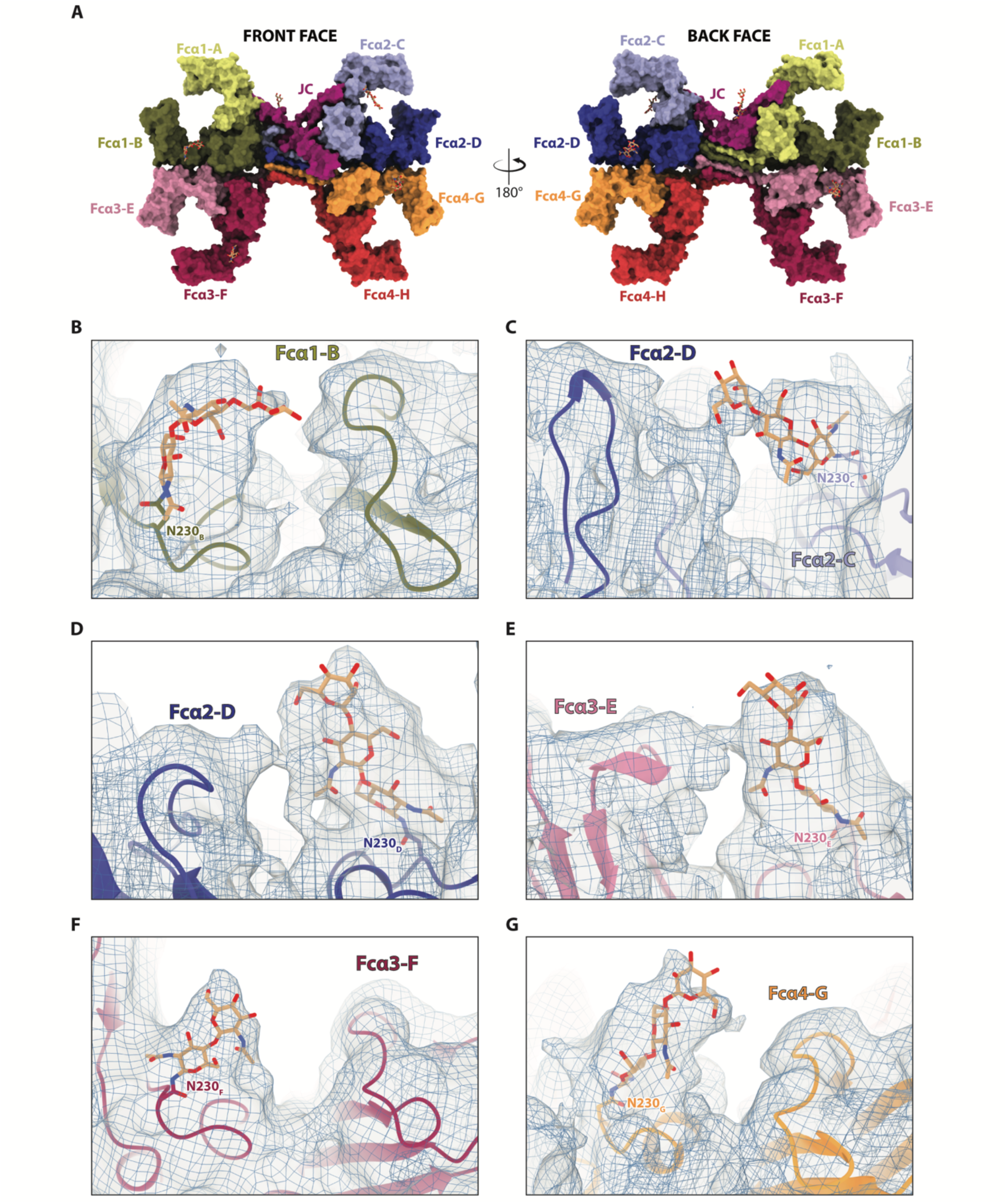
Heavy chain N-linked glycosylation modeled in *ap* JFcα cryo-EM map. (**A**) Front (left) and back (right) face views of *ap* JFcα with HCs and JC shown as surface representation and N-linked glycans shown as sticks and colored as in (Fig. 1A). (**B**-**G**) Focused view of N-linked glycans at Asn230 for chains Fcα1-B (B), Fcα2-C (C), Fcα2-D (D), Fcα3-E (E), Fcα3-F (F), and Fcα4-G (G). Each chain is shown as a cartoon representation with Asn230 and glycans shown as sticks. Cryo-EM density is shown using blue mesh and light grey transparent surface at the following thresholds: 0.08 (B and F), 0.055 (C), 0.06 (D), and 0.07 (E and G). _34_

**Fig. S4.**
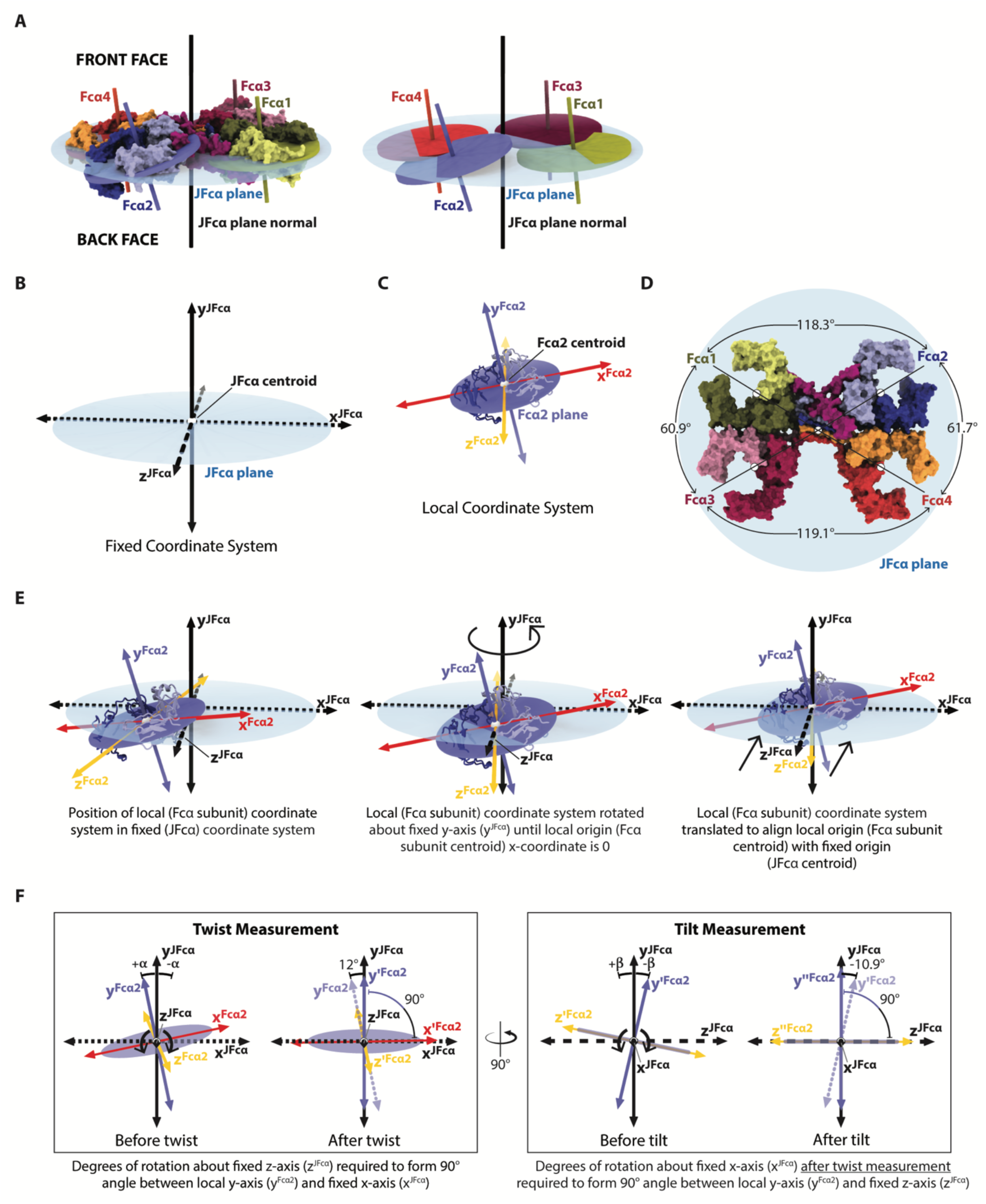
The unique geometric orientations between Fcα subunits in *ap* JFcα. (**A**) The *ap* JFcα structure shown with the JFcα plane (light blue ellipse) and with the central planes (solid ellipses) and associated normal (solid lines) for each Fcα. On the right, the *ap* JFcα structure is removed. (**B**) The fixed (JFcα) coordinate system with x^JFcα^-axis (dotted line), y^JFcα^-axis (solid line), and z^JFcα^-axis (dashed line). JFcα plane shown in light blue and JFcα centroid shown in white. (**C**) An example of a local (Fcα2) coordinate system with x^Fcα2^-axis (red line), y^Fcα2^-axis (blue violet line), and z^Fcα2^-axis (yellow line). The Fcα2 subunit is shown as a cartoon representation with the same coloring as in (A). The Fcα2 plane is shown as blue violet ellipse and the Fcα2 is centroid shown in white. (**D**) *ap* JFcα front face shown as molecular surface representation; subunits are colored as in (A) and labeled. JFcα plane and centroid are colored as in (B). Central angles (degrees) between adjacent Fcα subunits are indicated. (**E**) Origin alignment between fixed (JFcα) and local (Fcα2) coordinate systems prior to twist and tilt measurements. Fcα2 is shown as cartoon representation with same coloring as in (A). Fixed and local axes are distinguished and colored as in (B) and (C). Left, starting position of Fcα2 subunit. Center, position of local (Fcα2) coordinate system after rotation about fixed y-axis with local origin x-coordinate equal to 0. Right, position of local (Fcα2) coordinate system after translation to align local origin with fixed origin. (**F**) Twist (left) and tilt (right) measurements for Fcα2 subunit. Axes for the local coordinate system after the twist and tilt rotations are indicated with a prime or double prime marking, respectively.

**Fig. S5.**
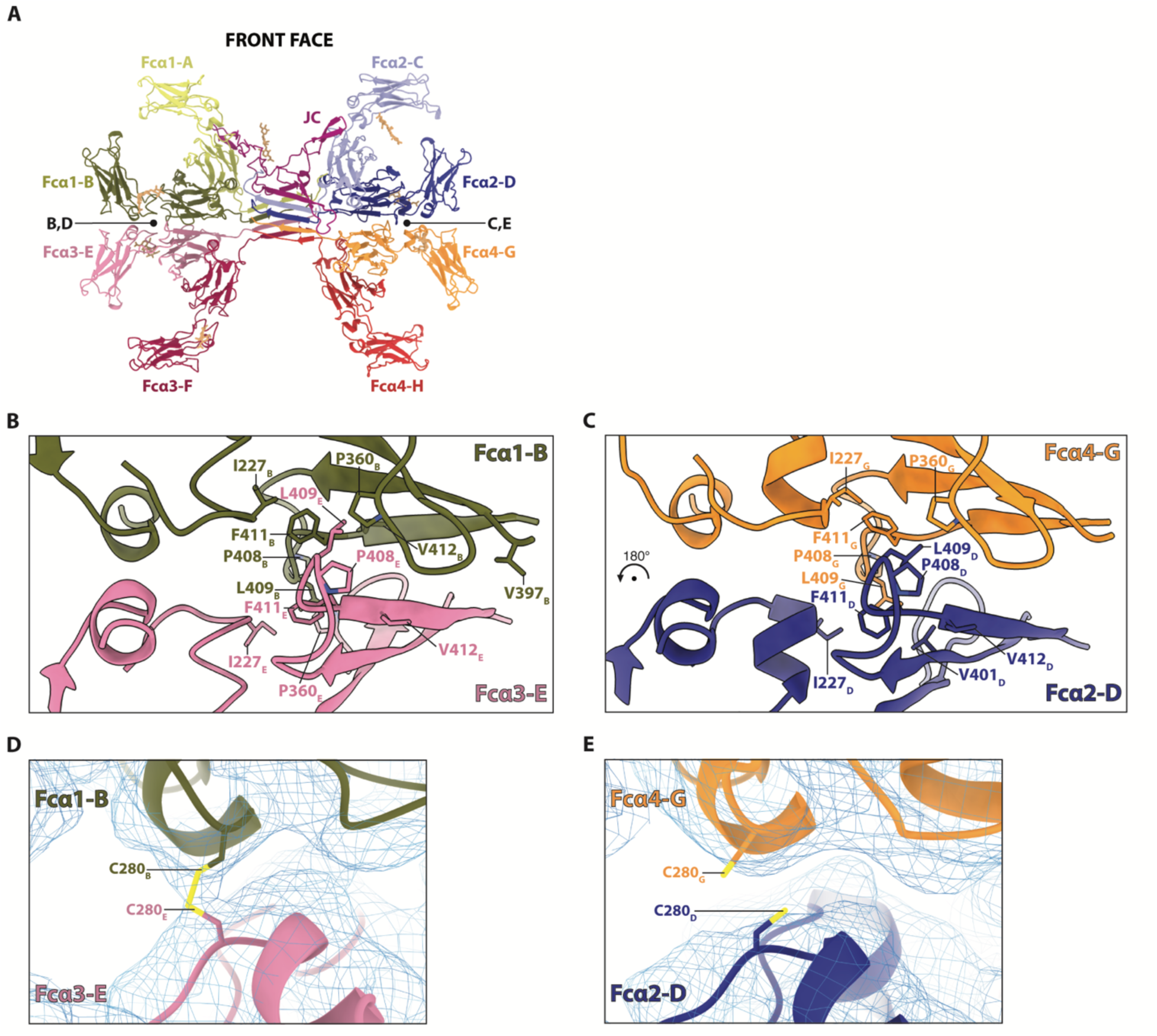
Molecular interactions at Fc-Fc interfaces in *ap* JFcα. (**A**) Front face view of *ap* JFcα shown as cartoon with glycans shown as sticks. Colors are same as in (Fig. 1A). Black lines and letters indicate focused regions for subsequent panels. (**B**) Fcα1-Fcα3 and (**C**) Fcα2-Fcα4 interfaces shown as cartoon representations with hydrophobic residues participating in intra-Fc interactions labeled and shown as sticks. Chains and residues are colored as in (Fig. 1A). (**D-E**) Focused and magnified view of Fcα1-Fcα3 (D) and Fcα2-Fcα4 (E) interfaces where Cys280 residues are shown as sticks. Cryo-EM map density is also shown and carved to a threshold of 0.12.

**Fig. S6.**
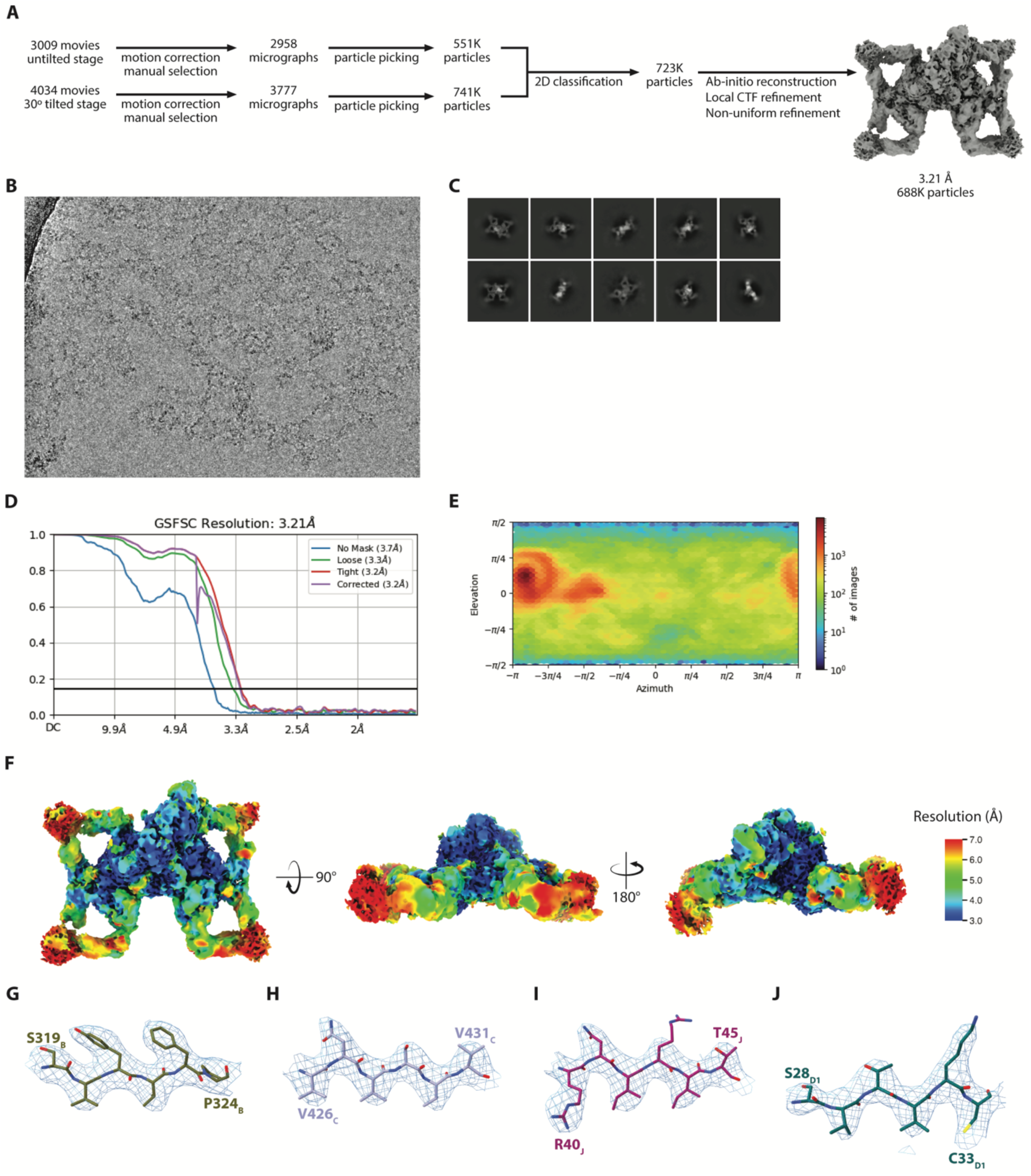
Cryo-EM data collection and CryoSPARC processing pipeline for *ap* SJFcα. (**A**) Schematic summary of the *ap* SJFcα cryo-EM data processing pipeline in CryoSPARC. (**B**) Representative micrograph of *ap* SJFcα. (**C**) Representative 2D class averages. (**D**) The FSC curve for the final reconstruction with reported resolution at FSC=0.143 shown by the (color) horizontal line. (**E**) Local resolution map calculated by CryoSPARC and colored in Chimera. (**F**) Viewing direction distribution plot of particles used in final refinement. (**G-J**) Cryo-EM map density surrounding residues 319 to 324 in Fcα1-A Cα4 (G), residues 426 to 431 in Tp_C_ (H), residues 40 to 45 in JC_core_ (I), and residues 28 to 33 in SC (J), each contoured to threshold of 0.13 and carved at 2 Å.

**Fig. S7.**
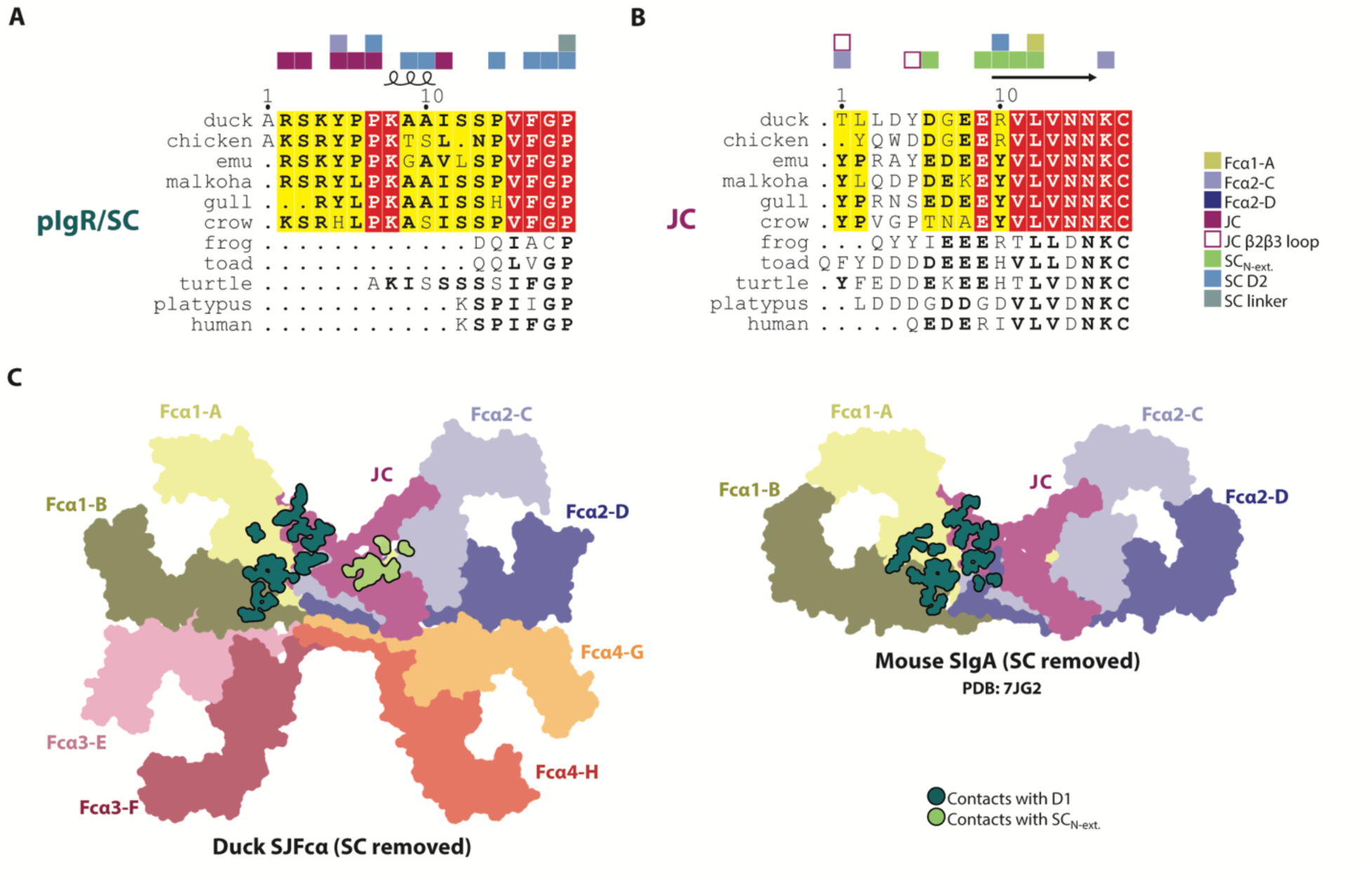
Comparison of SC D1 binding sites in *ap* SJFcα and mouse SIgA. (**A**) Sequence alignment of pIgR/SC N-terminal sequences from representative vertebrate species with duck SC_N-ext._ contacts indicated above and colored according to the key. (**B**) Sequence alignment of JC N-terminal sequences in representative vertebrate species with contacts of duck JC_N-ext._ contacts indicated above and colored according to the key. (**C**) Left, flat surface representation of *ap* SJFcα with SC removed. Right, flat surface representation of mouse SIgA (PDB: 7JG2) with SC removed. Surfaces colored uniquely and labeled. HC and JC residues contacting SC D1 or the SC_N-ext._ are colored dark teal or lime green, respectively.

**Fig. S8.**
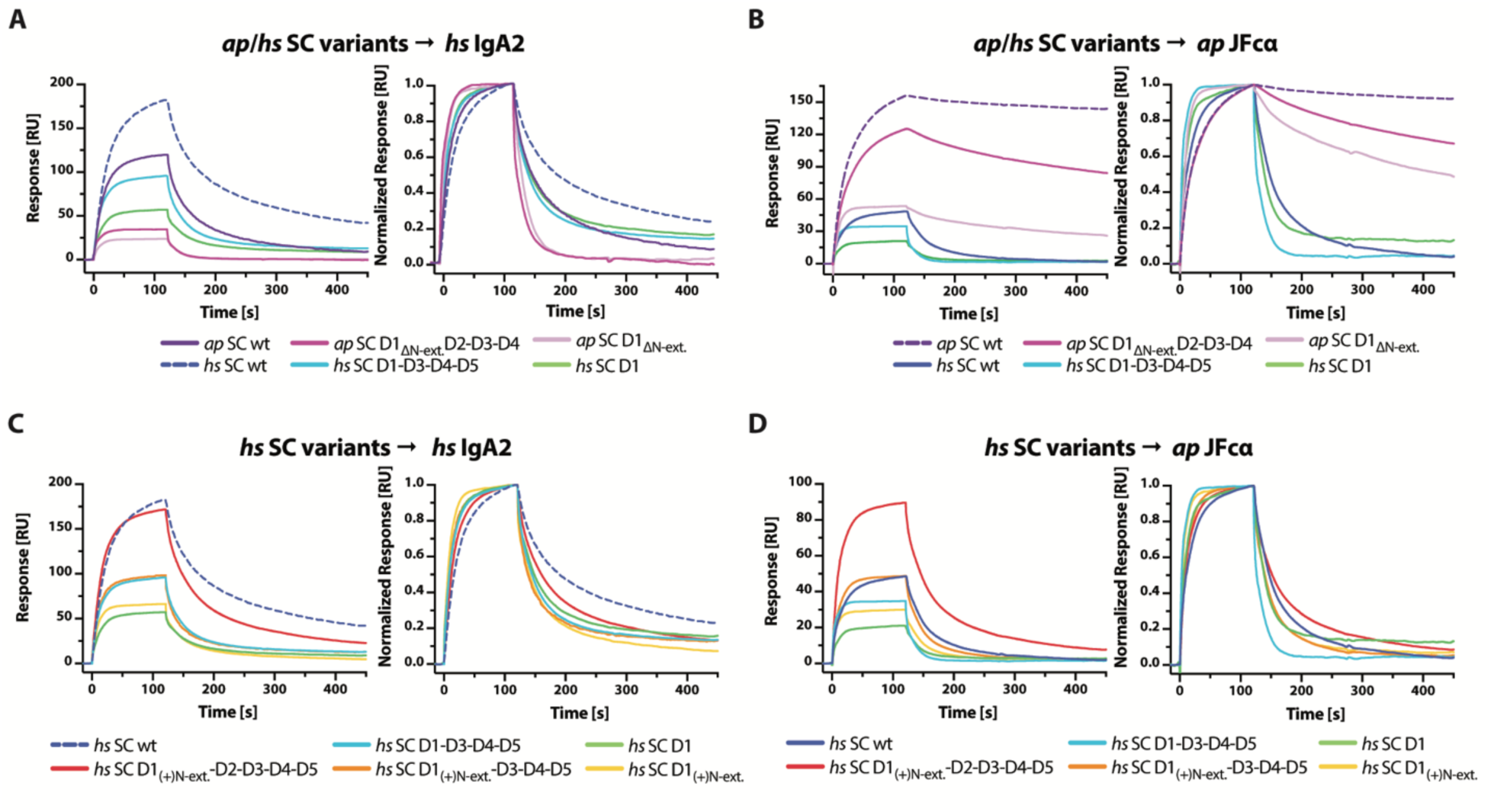
Duck and human SC variant binding to cognate and non-cognate ligands. (**A-D**) SPR sensorgrams showing the response (left) and normalized response (right) of 64nM *ap* SC and *hs* SC variant analytes binding to either *hs* IgA2 (A) or *ap* JFcα (B), as well as *hs* SC and chimeric *hs* SC variants binding to either *hs* IgA2 (C) or *ap* JFcα (D). Each analyte response is colored according to the key at the bottom of the panel and dashed lines indicate cognate interactions between unmutated variants. Results are consistent with three or more replicate experiments. Sensorgrams for the full analyte concentration series of each experiment are provided in fig. S9, 11-12.

**Fig. S9.**
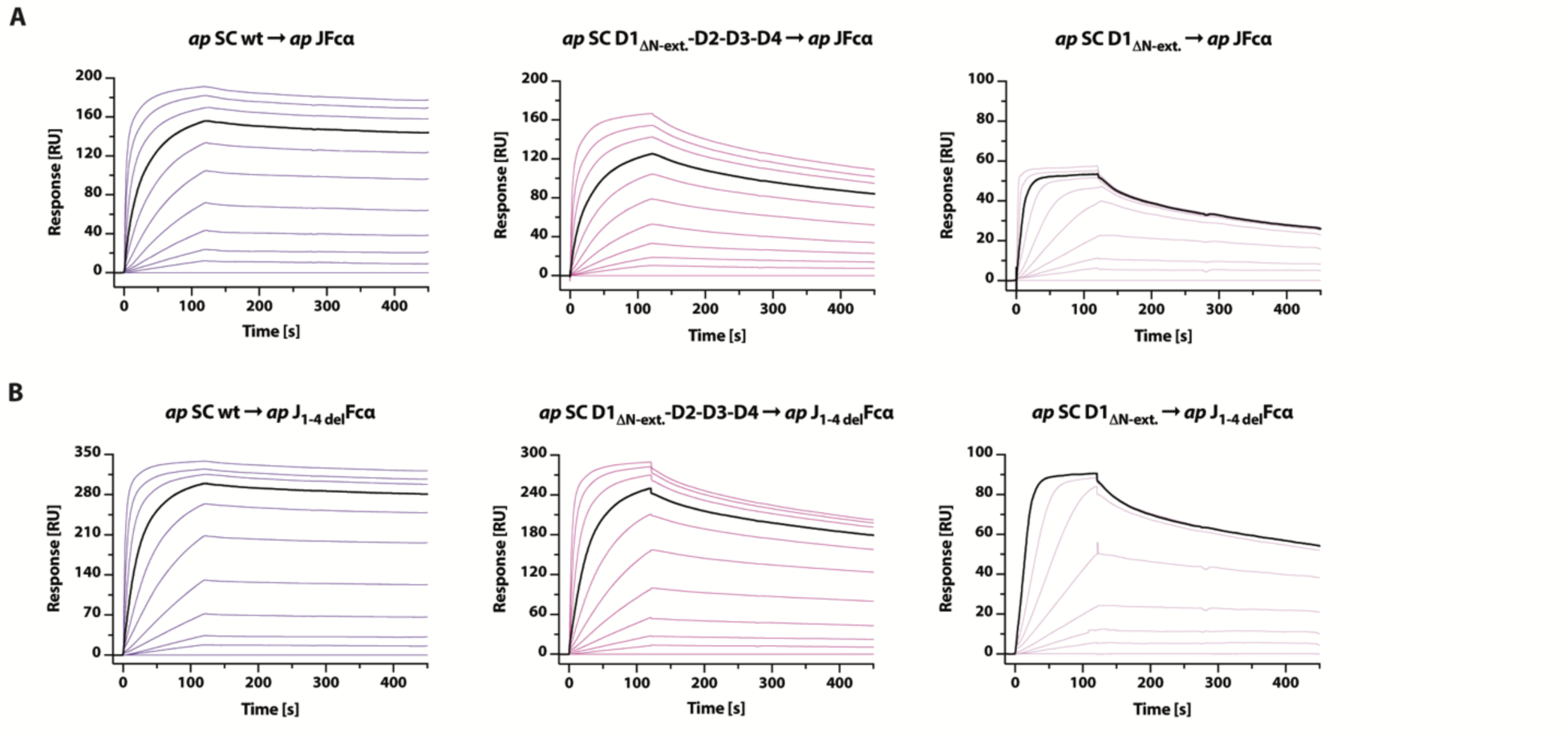
Duck SC variants binding to *ap* JFcα. (**A-B**) SPR sensorgrams showing responses for entire two-fold dilution series of *ap* SC variants binding to *ap* JFcα (A) and *ap* J_1-4del_Fcα (B). The maximum concentration used was 512 nM for all analytes, except for *ap* SC D1_ΔN-ext._ binding to *ap* J_1-4del_Fcα, which had a maximum concentration of 64nM. Sensorgrams are colored to match corresponding curves in concentration-matched figures (Fig. 8; fig. S8), except for responses at 64 nM, which are colored black. Sensorgrams shown are equivalent to those obtained from replicates (not shown).

**Fig. S10.**
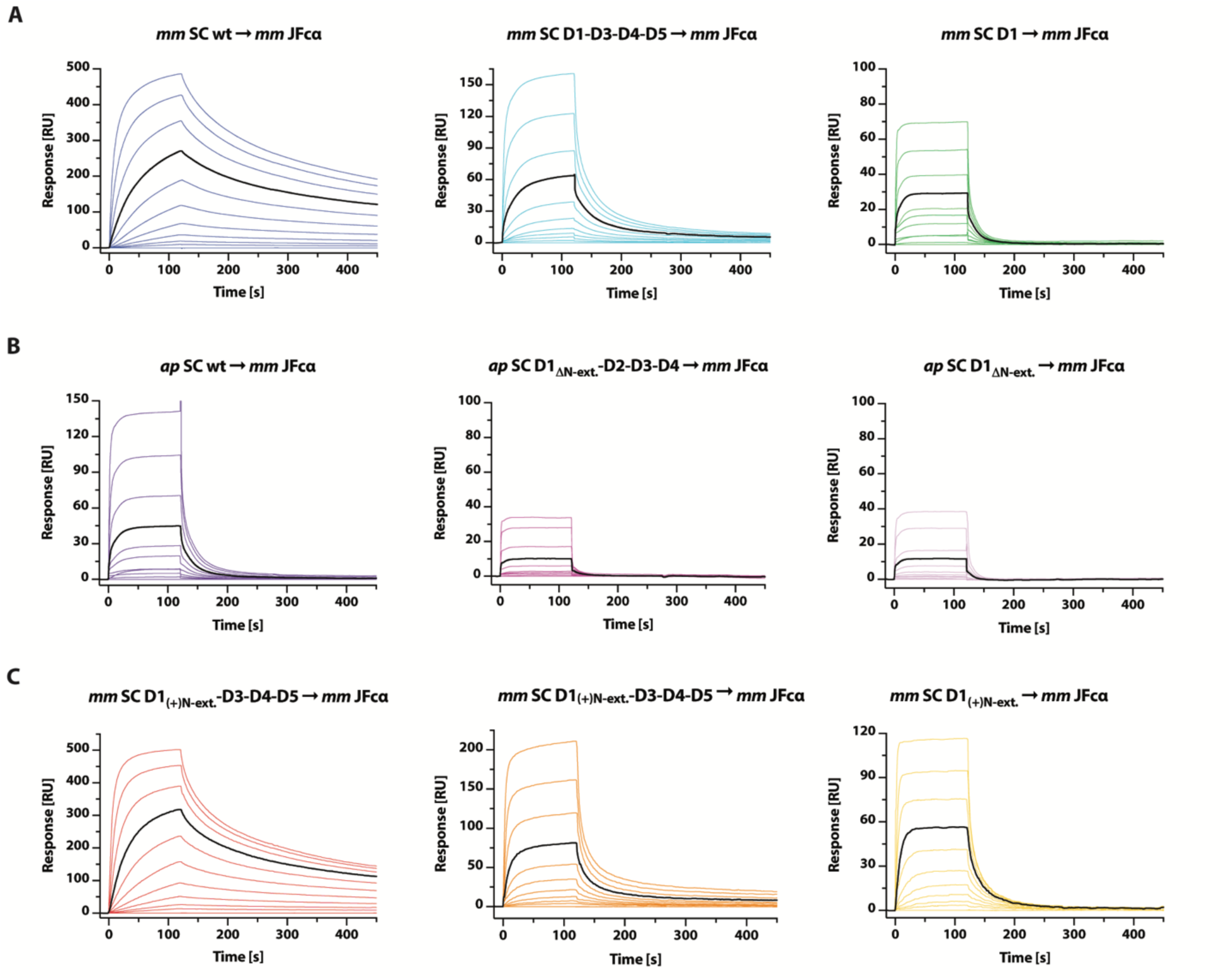
Mouse and duck SC variants binding to mouse JFcα. (**A-C**) SPR sensorgrams showing responses for entire two-fold dilution series of *mm* SC (A), *ap* SC (B), or chimeric *mm* SC (C) variants binding to *mm* JFcα. The maximum concentration used was 512 nM for all analytes. Sensorgrams are colored to match corresponding curves in concentration-matched figures (Fig. 8), except for responses at 64 nM which are colored black. Sensorgrams shown are equivalent to those obtained from replicates (not shown).

**Fig. S11.**
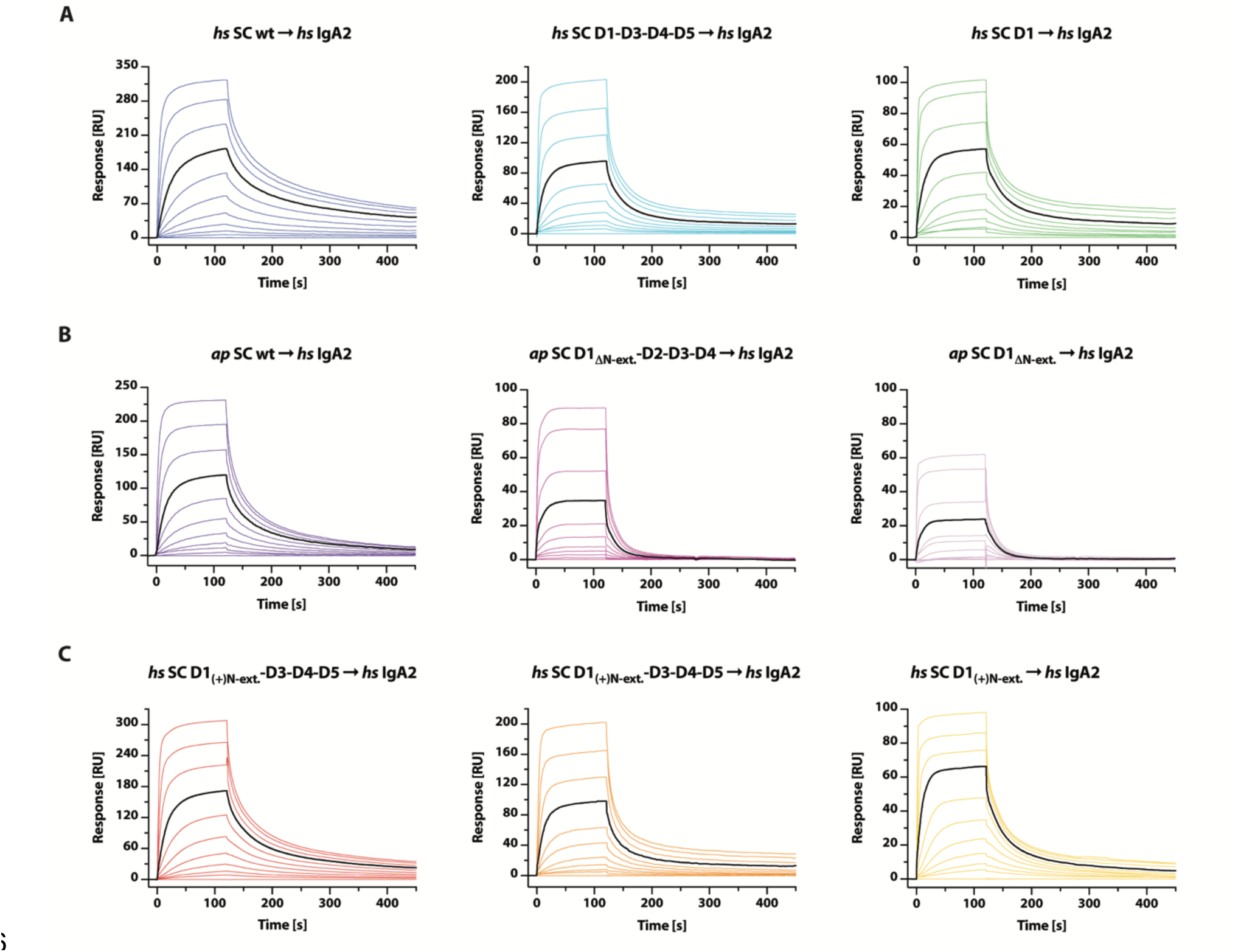
Human and duck SC variants binding to human IgA2. (**A-C**) SPR sensorgrams showing responses for entire two-fold dilution series of *hs* SC (A), *ap* SC (B), or chimeric *hs* SC (C) variants binding to *hs* IgA2. The maximum concentration used was 512 nM for all analytes. Sensorgrams are colored to match corresponding curves in concentration-matched figures (fig. S8), except for responses at 64 nM, which are colored black. Sensorgrams shown are equivalent to those obtained from replicates (not shown).

**Fig. S12.**
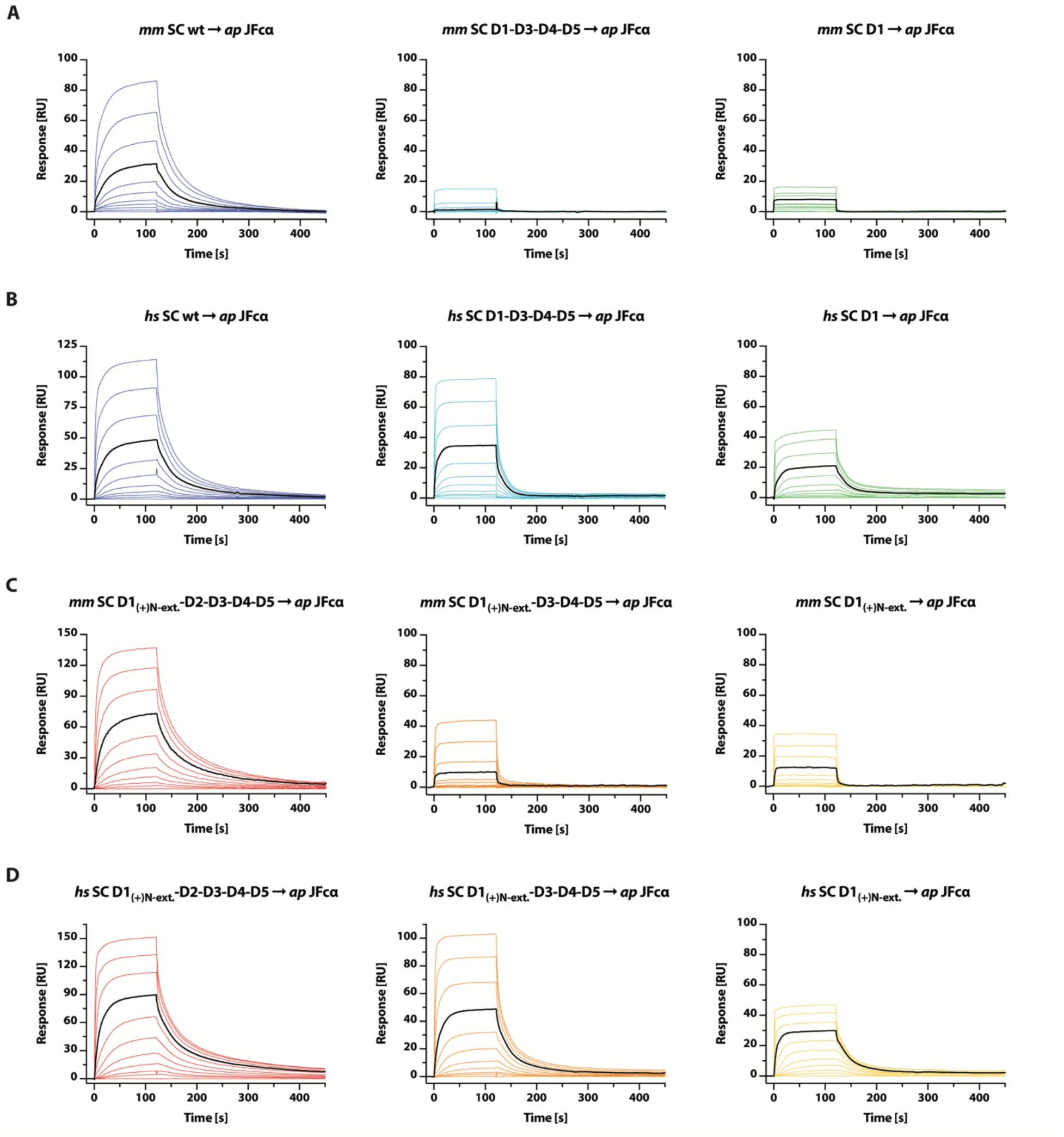
Mouse and human SC variants binding to *ap* JFcα. (**A-D**) SPR sensorgrams showing responses for entire two-fold dilution series of *mm* SC (A), *hs* SC (B), chimeric *mm* SC (C), and chimeric *hs* SC (D) variants binding to *ap* JFcα. The maximum concentration used was 512 nM for all analytes. Sensorgrams are colored to match corresponding curves in concentration-matched figures (Fig. 8; fig. S8), except for responses at 64 nM, which are colored black. Sensorgrams shown are equivalent to those obtained from replicates (not shown).

**Table S1.** Cryo-EM data collection and model refinement statistics.

|  | **ap JFα**<br>**EMD 47901**<br>**PDB 9EC7** | **ap SJFα**<br>**EMD 47900**<br>**PDB 9EC6** |
| --- | --- | --- |
| **Data collection** |  |  |
| **Voltage (kV)** | 300 | 300 |
| **Nominal magnification** | 81,000x | 105,000x |
| **Total dose (electrons/Å<sup>2</sup>)** | 57.79 | 71.3 |
| **Total exposure time (s)** | 3.21 | 2.41 |
| **Raw movie pixel size (Å/pixel)** | 0.527 | 0.822 |
| **Defocus range (μm)** | 0.1 – 2.7 | 0.1 – 2.7 |
| **Frames per movie** | 40 | 60 |
| **Automation software, untilted/tilted** | EPU / Leginon | EPU / SerialEM |
| **Data processing** |  |  |
| **Movies collected, untilted/tilted** | 2544 / 3015 | 3009 / 4034 |
| **Initial/final particles** | 10603K / 320667 | 1392K / 688138 |
| **Map resolution (Å)** | 3.76 | 3.21 |
| **FSC threshold** | 0.143 | 0.143 |
| **Density and fit** |  |  |
| **Model vs. data (CC)** |  |  |
| **Mask** | 0.75 | 0.78 |
| **Box** | 0.80 | 0.84 |
| **Peaks** | 0.64 | 0.69 |
| **Volume** | 0.75 | 0.78 |
| **Model composition** |  |  |
| **Total atoms** | 29201 | 36076 |
| **Non-hydrogen atoms** | 14758 | 18277 |
| **Protein residues** | 1917 | 2365 |
| **Ligands** | BMA: 6; NAG:14 | BMA: 4; NAG: 17 |
| **Bond RMSD** |  |  |
| **Lengths (Å)** | 0.004 | 0.003 |
| **Angles (°)** | 0.966 | 0.459 |
| **MolProbity score** | 1.67 | 1.64 |
| **Clash score** | 4.94 | 4.99 |
| **Ramachandran plot (%)** |  |  |
| **Outliers** | 0.00 | 0.00 |
| **Allowed** | 6.16 | 5.54 |
| **Favored** | 93.84 | 94.46 |
| **Rotamer outliers (%)** | 0.00 | 0.00 |
| **Cβ outliers (%)** | 0.00 | 0.00 |
| **Cis proline** | 0.0 | 0.0 |
| **Twisted proline** | 0.0 | 0.0 |
| **CaBLAM outliers (%)** | 4.25 | 4.65 |

**Table S2.**
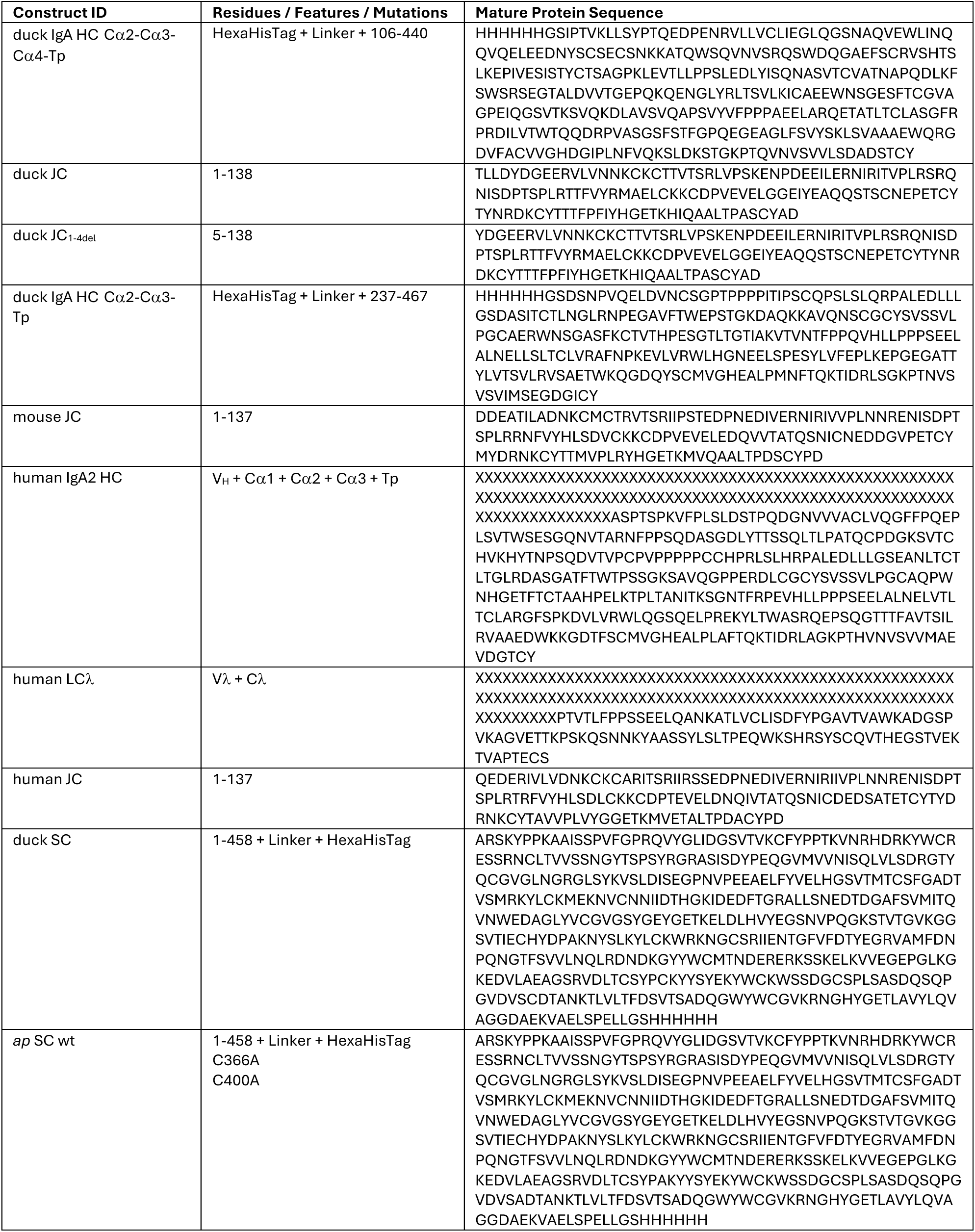
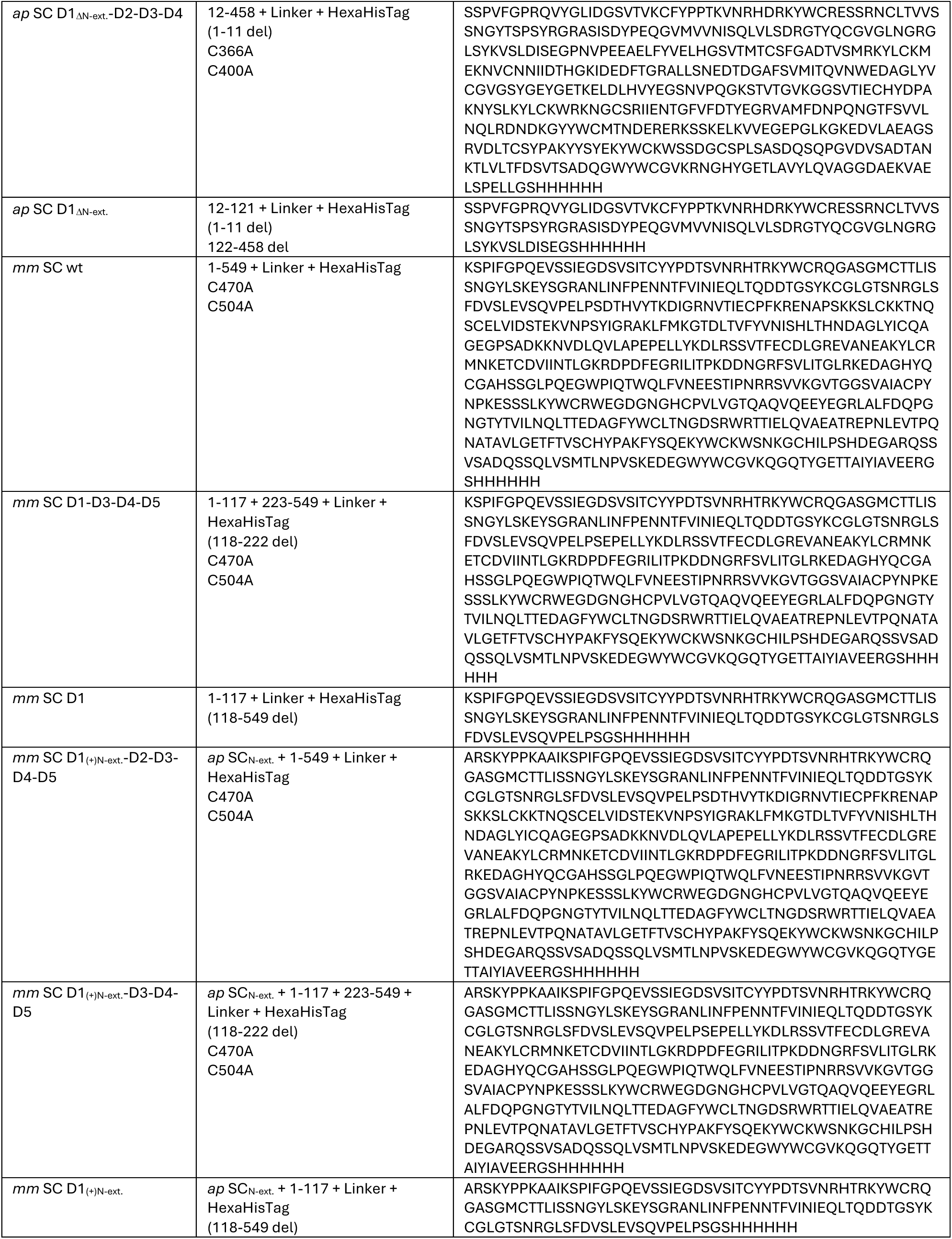
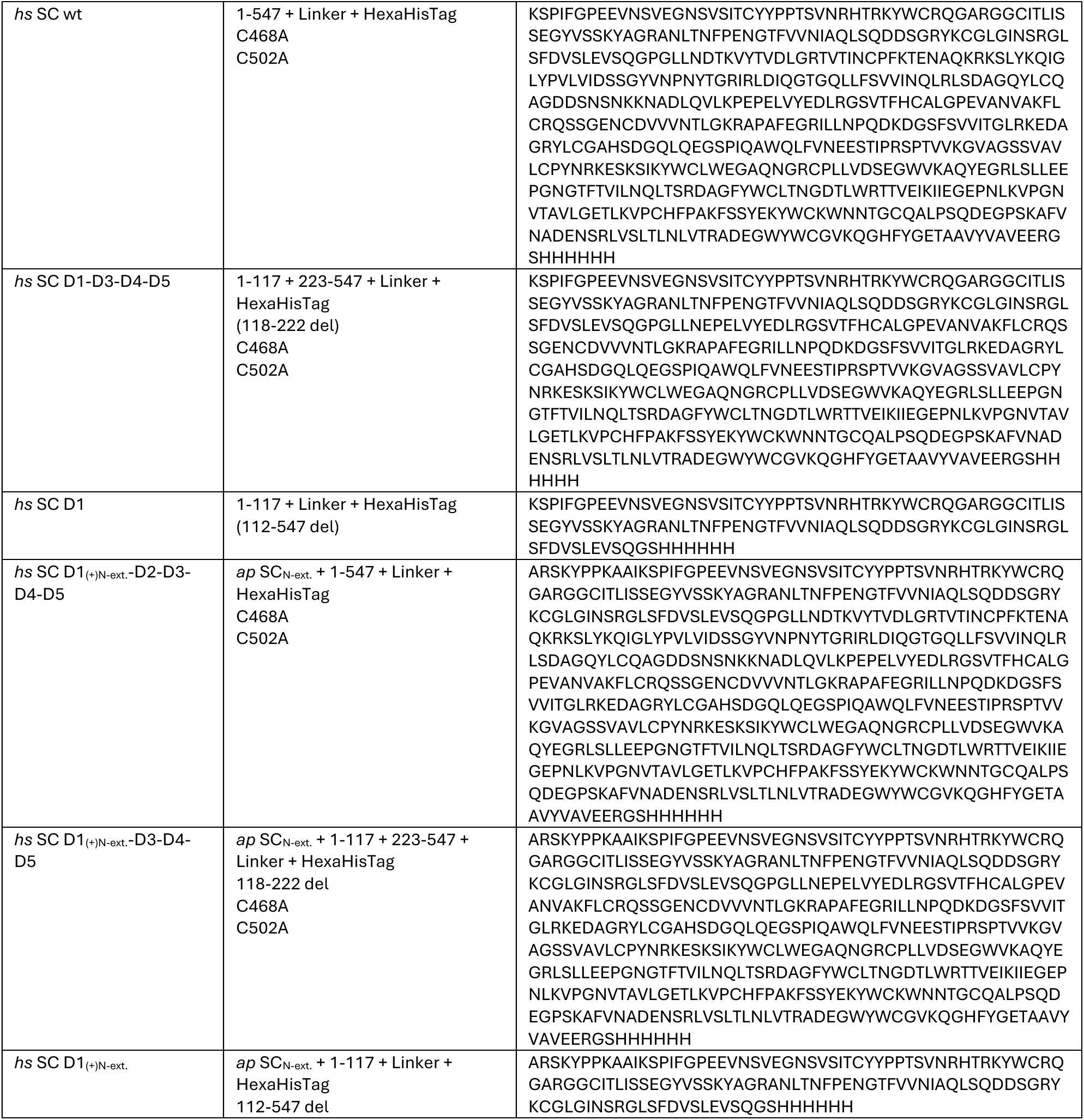
Expression constructs used in this study.

